# Disinhibitory signaling enables flexible coding of top-down information

**DOI:** 10.1101/2023.10.17.562828

**Authors:** Tomas Gallo Aquino, Robert Kim, Nuttida Rungratsameetaweemana

## Abstract

Recent studies have proposed employing biologically plausible recurrent neural networks (RNNs) to investigate flexible decision-making in the brain. However, the mechanisms underlying the integration of bottom-up sensory inputs and temporally varying top-down factors (such as task instructions and selective attention) remain poorly understood, both within the context of these models and the brain. To address this knowledge gap, we trained biologically inspired RNNs on complex cognitive tasks that require adaptive integration of these factors. Through comprehensive analyses of RNNs and neural activity from mouse primary visual cortex, we show that sensory neurons in low-level areas possess the remarkable ability to multiplex and dynamically combine both bottom-up and top-down information via local inhibitory-to-inhibitory connections. Our results shed light on the role of disinhibitory circuits in the intricate interplay between bottom-up and top-down factors to enable flexible decision processes.

## Introduction

Successfully navigating complex environments requires neural systems to efficiently encode and process incoming sensory inputs alongside contextual cues and task goals [1–3]. This capacity to integrate across different sources of information allows the brain to dynamically convert identical sensory inputs into diverse behavior in a context-dependent manner. Sensory inputs can be understood as bottom-up signals with regards to their encoding within lower cortical areas. In contrast, top-down factors refer to information regarding behavioral relevance of each stimulus, prior expectations about sensory statistics, and task contexts which provide guidelines for how the encoded sensory information should be interpreted and mapped onto behavior [4–8].

The ability to rapidly adapt to temporally varying top-down factors is a hallmark of cognitive flexibility and has been primarily attributed to the frontoparietal networks and medial temporal lobes [9–15]. However, the extent to which lower cortical areas contribute to this adaptive process is not well understood. Past works have shown that responses within sensory areas could be modulated by top-down signals such as selective attention [16–25], and that non-sensory information such as memory content can be decoded from activity within sensory areas [26–29]. Yet, it remains poorly understood how these neural codes complement computations that take place in higher cortical areas. This knowledge gap highlights the need for deeper insight into the computational principles and circuit mechanisms that enable flexible information processing for adaptive behavior.

In nature, where identical stimuli elicit diverse responses, inhibition may be a key mechanism in averting conflicting behaviors. This ensures the selection of appropriate sensorimotor mapping, aligning behavioral outcomes with the current task contexts rather than being driven solely by the stimuli. This account of inhibition as a modulatory mechanism, crucial for shaping task-optimized behavior, has been supported by past studies demonstrating the role of inhibition for the integration of top-down factors within the cortical hierarchy [17, 30–32]. In addition, discoveries of specialized sensory microcircuits that rely on inhibitory-to-inhibitory connections among local interneurons further highlight the importance of inhibitory function in flexible information processing [33–36]. Notably, disinhibitory circuits have been proposed as key in facilitating top-down modulation of sensory coding [37–40], highlighting the plastic nature of neural codes within lower cortical areas. This account underscores the flexibility of neural codes along the cortical hierarchy and suggests a potential mechanism for sensory multiplexing [41, 42].

Models based on recurrent neural networks (RNNs) of continuous-variable firing rate units have been widely used to provide computational explanations of experimental findings and to investigate neural correlates of cognitive functions, including sensory discrimination and working memory [9, 43–49]. However, these studies primarily examine scenarios in with task goals and contextual cues remain fixed over time. This results in the models basing their responses largely on bottom-up processes such that the generated behaviors are mainly governed by the encoding of sensory inputs.

This approach may not fully capture the dynamics of the natural environment, where top-down information about current task demands is consistently present, exerting continuous influence on how we optimize the conversion of sensory inputs into appropriate behaviors. Importantly, these top-down factors, including selective attention, task contexts, and prior knowledge, can exhibit distinct temporal structures, imposing varied but complementary guiding rules on how sensory input should be processed. This results in temporally rich and highly dynamic decision-making, a phenomenon that has proven challenging to investigate in artificial neural network models, including RNNs. Neglecting to account for the nuanced dynamics of top-down factors and their temporal influence limits our understanding of how neural systems efficiently integrate bottom-up and top-down signals to consistently produce optimal behavior. Additionally, we can gain better insights into this dynamic process through modifying the architecture of past models to incorporate several fundamental properties inherent in biological neural networks. These constraints, such as heterogenous neuronal timescales, cortical hierarchies, and the balance between inhibitory and excitatory connections, can offer valuable insights into the functionality of neural networks and the types of computations they can perform [50–53].

To address these knowledge gaps, we designed dynamic decision-making tasks that require seamless integration of bottom-up sensory inputs with top-down factors. These tasks included diverse top-down signaling with varying temporal dynamics as well as orthogonal manipulation of bottom-up and top-down signals. By training biologically plausible RNNs on these tasks and parsing the trained networks, we show that disinhibitory circuit motifs governed by inhibitory-to-inhibitory connections are critical for the intricate interplay between the two essential components of decision processes. By extending our model to incorporate the functional cortical hierarchy, we further demonstrate that feedback signaling from higher-level areas and selective local inhibition enable lower-level areas to differentially process identical sensory inputs in a context-dependent manner. Moreover, we tested our model predictions using publicly available data recorded from the mouse primary visual cortex [54] and confirmed that local inhibitory-to-inhibitory connections within the visual cortex play an integral role in encoding top-down information.

Our results offer mechanistic insights into the dynamic integration of top-down and bottom-up signals during flexible information processing in a biological context. Within this comprehensive theoretical framework, we elucidate the computational principles employed by RNN models to flexibly utilize distinct top-down signals for guiding the analysis of sensory input, thereby optimizing behavior.

## Results

### Recurrent neural network (RNN) model and dynamic decision-making tasks

To understand the cortical computations required for flexibly and optimally integrating bottom-up and top-down information, we employed a biologically plausible recurrent neural network (RNN) model consisted of continuous-variable firing units (model schematic shown in Fig. 1a right; see *Methods*) and trained the model to perform novel dynamic decision-making tasks (Fig. 1a,b). We utilized a gradient descent approach to train the RNN model, optimizing the recurrent connectivity weights, readout output weights, and synaptic decay time constants (see *Methods* for details). In addition, Dale’s principle was enforced using the weight parametrization method introduced in [43].

**Fig. 1.**
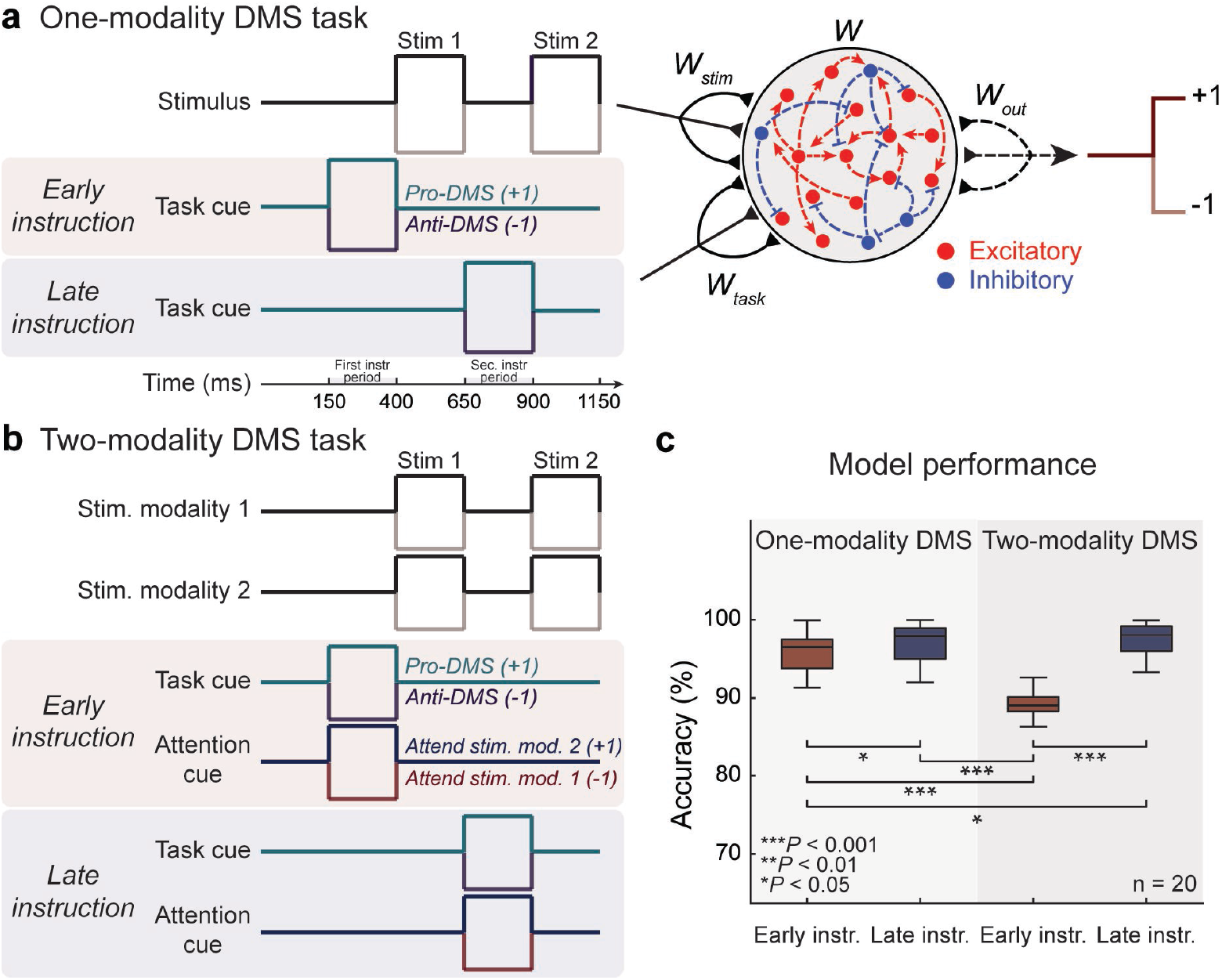
Dynamic decision-making tasks and model schematic. **a.** Schematic of a one-modality DMS task with sequential stimuli separated by a delay period. A task cue, indicating the stimulus-response mapping, is presented either before (*early instruction*) or after stimulus 1 (*late instruction*). The recurrent neural networks (RNNs) generate an output of +1 or -1 for matched and mismatched trials, respectively (*pro-DMS*; and vice versa for *anti-DMS*). The schematic of the RNN model shown on the right. **b.** Schematic of a two-modality DMS task where each stimulus comprises two modalities. In addition to the task cue, an attention cue is presented on each trial, indicating the modality on which the RNNs need to perform the DMS task. The task and attention cues are presented either before (*early instruction*) or after stimulus 1 (*late instruction*). The trial timing for both one- and two-modality DMS tasks is depicted in **a**. **c.** Testing performance of the trained RNNs on the DMS tasks. The average accuracy of the trained RNNs is presented, with statistical comparisons computed using two-sided Wilcoxon rank-sum tests. Boxplot: central lines, median; bottom and top edges, lower and upper quartiles; whiskers, 1.5 × interquartile range; outliers are not plotted. \**p <* 0.05; \*\**p <* 0.01; \*\*\**p <* 0.001

These RNNs were employed to execute flexible decision-making tasks designed to achieve several distinct objectives. The first aim was to investigate how identical sensory inputs might be processed differently depending on distinct top-down inputs. These tasks also allowed for the independent manipulation of both bottom-up and top-down inputs. Lastly, the task incorporated separate stimulus modalities, each providing independent sources of top-down information. To meet these criteria in task design, we modified the classic delayed match-to-sample (DMS) task, commonly used in experimental studies [55–59], to develop one-modality DMS and two-modality DMS tasks.

For the one-modality DMS task (Fig. 1a), the RNN model received one stream of input stimulus signal (bottom-up) containing two sequential stimuli separated by a delay period. Each stimulus was set to either +1 or -1. The model additionally received a task cue signal (top-down), indicating whether the network should employ the pro- or anti-DMS stimulus-response mapping. For the pro-DMS task (a task cue of +1 during the instruction period), the network was required to generate an output of +1 if the signs of the two stimuli matched, and -1 otherwise. For the anti-DMS (a task cue of -1 during the instruction period), the network was required to generate a negative output if the signs of the two stimuli were the same, and a positive output otherwise. The task cue was given either before (*early instruction*; 150-400 ms) or after the first stimulus (*late instruction*; 650-900 ms; Fig. 1a).

Critically, in nature, not all aspects of sensory input are relevant to the task at hand (i.e., one-modality DMS task). To effectively navigate this, organisms rely on additional top-down signals, such as selective attention, alongside direct task signals to optimize decisions. To study this more complex scenario, we developed the two-modality DMS task by incorporating two streams of input stimulus signals (bottom-up; modality 1 and modality 2) and an attention cue (top-down) which instructs the model which stimulus modality is behaviorally relevant on a given trial and thus should be attended to (Fig. 1b; see *Methods*). An attention signal of -1 during the instruction period informs the model to focus on modality 1 while ignoring modality 2, and vice versa for a +1 attention cue signal. The task cue manipulation is identical to that used in the one-modality DMS task. Similar to the one-modality task, the timing of the instruction period was manipulated such that task and attention cues were delivered either before (*early instruction*) or after the presentation of the first stimulus (*late instruction*).

For each decision task (one-modality DMS and two-modality DMS), we trained 20 RNNs to perform the task with high accuracy (*>* 95%; see *Methods*). For the one-modality DMS, the models performed the task accurately regardless of the timing of the instruction window (Fig. 1c). The model performance on the two-modality DMS was lower when the top-down cues were provided early (*p <* 0.001, two-sided Wilcoxon rank-sum test Fig. 1c). This can be attributed to the model’s requirement to retain both the task and attention signals for a longer duration in the early instruction condition.

### Dynamic encoding of top-down information by RNNs

After training RNNs to perform the dynamic decision tasks, we next investigated the effects of the top-down factors (task and attention cues) on the network dynamics. To visualize RNN dynamics in response to distinct external inputs of both bottom-up and top-down signals under various conditions, we plotted the kinetic energy landscape (see *Methods*) of a representative RNN. This was achieved by using two principal components (Fig. 2a,b; see *Methods*). The kinetic energy landscape provides a useful guideline for understanding RNNs as dynamical systems. It offers a visual summary that illustrates how external inputs shape the structure of fixed points and “slow” points (local minima in the energy landscape) as well as neural trajectories within the state space [60–62]. Therefore, seeking to identify these fixed and “slow” points in the two-dimensional energy landscape enabled us to gain insight into how the task and attention signals influenced the RNN network dynamics.

**Fig. 2.**
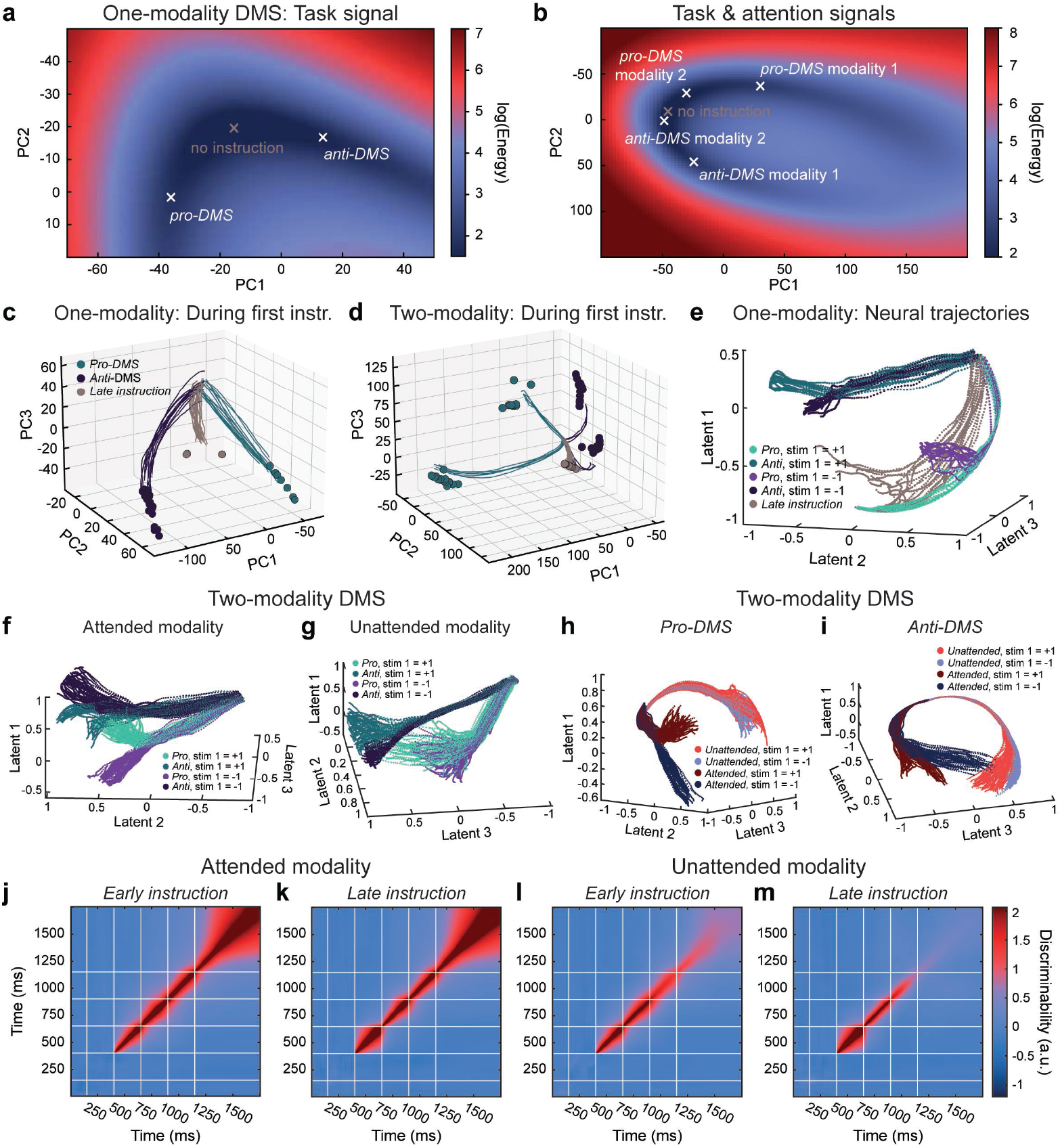
Flexible incorporation of task and attention signals. **a.** Heat map of kinetic energy in an example RNN performing the one-modality DMS task during the first instruction period (150-400 ms; Fig. 1a). Background heat maps belong to the late instruction condition. The middle cross indicates the local energy minimum during the first instruction period for the late instruction condition (*no instruction*), while the left and right crosses indicate the new local energy minima for the pro- and anti-DMS conditions, respectively. **b.** Same as **a**, for an example RNN performing the two-modality DMS task. Legend: pro-task modality 1, pro-DMS and first modality attended; pro-task modality 2, pro-DMS and second modality attended; anti-task modality 1, anti-DMS and first modality attended; anti-task modality 2, anti-DMS and second modality attended. **c.** Individual trial one-modality RNN trajectories in principal component space during the first instruction period. Principal component projections of fixed points are included. Trial top-down cue conditions are divided into pro-DMS (teal), anti-DMS (dark blue) or no instructions (gray). **d.** Same as **c**, for a two-modality RNN. **e.** Latent space one-modality RNN trajectories during the furst instruction and first stimulus periods (150-650 ms; Fig. 1a) for all combinations of top-down cues and stimulus identities. **f.** Same as **e**, for a two-modality RNN, highlighting the attended stimulus modality. **g.** Same as **e**, for a two-modality network, highlighting the unattended stimulus modality. **h.** Latent space two-modality RNN trajectories for pro-DMS trials, splitting trajectories by the identity of attended versus unattended stimuli. **i.** Same as **h**, for anti-DMS trials. **j-k.** Temporal discriminability for the first stimulus identity of the attended modality when the attention signal is given early (**j**) or late (**k**). The task signal was fixed to anti-DMS. **l-m.** Temporal discriminability for the first stimulus identity of the unattended modality when the attention signal is given early (**l**) or late (**m**). The task cue was fixed to anti-DMS.

We hypothesized that the presence of top-down cues would condition network dynamics even before the arrival of the bottom-up sensory inputs. In addition, we postulated that opposing cues would correspondingly lead to separable shifts in fixed point structure, which we first visualized through the energy landscape of the network during the first instruction period (150-400 ms, see Fig. 1a). For the one-modality DMS task, the energy landscape of the example RNN model displayed an area with an inverted U-shape where the kinetic energy was notably low (darker blue area in Fig. 2a). The local minimum of the landscape stayed around the center of this U-shaped profile during the period before the first stimulus presentation when the task cue was not yet presented (*no instruction*). However, when the task cue was presented early (*early instruction*), the local minimum shifted to the left and to the right for the pro- and anti-DMS condition, respectively (Fig. 2a).

For the two-modality DMS task, the energy landscape again revealed a U-shaped region characterized by a low-energy profile (dark blue in Fig. 2b). Consistent with what we observed in the one-modality task, the local minimum was situated at the center of the low energy structure during the first instruction period for late instruction trials, where no task or attention cues were yet presented (*no instruction*). Analyzing different combinations of the task and attention cue signals displayed distinct local minima points along the low-energy profile. For instance, the local minima corresponding to the pro-DMS signal were positioned to the right of the no-instruction minima. Along this pro-DMS “arm” of the energy trajectory, two local minima were observed: one corresponding to the modality-one attention signal and the other to the modality-two attention signal. These results suggest that network dynamics shift as a function of the combination of top-down attention and task signals, in a dissociable manner even before the sensory inputs arrive.

Subsequently, we performed a quantitative fixed point search analysis [60] for different values of external input to the network during the first instruction period (150-400 ms, Fig. 1a). During this time, the network could receive either a pro-DMS, an anti-DMS, or no task cue (i.e., *late instruction* trials). We plotted network trajectories as well as fixed points discovered for each of these conditions in the principal component space, for the representative RNN performing the one-modality (Fig. 2c) and the two-modality DMS tasks (Fig. 2d). We observed that the presence and content of top-down signals given to the network not only altered its trajectories in the state space but also affected its fixed point structure. The network performing the two-modality DMS task exhibited a branching pattern that reflected symmetrical top-down coding across attended modalities for each task cue condition (anti- vs. pro-DMS).

To further examine network dynamics as a function of distinct top-down and bottom-up information, we performed a manifold discovery analysis, utilizing the CEBRA method [63]. This approach enables latent structure discovery and has the advantage of conditioning latent structure discovery over variables of interest. Here, we performed an unsupervised analysis with regards to all task covariates except for time. This ensured that adjacent time points were mapped to closer points in latent space and that smooth latent dynamics were obtained. This method allowed us to determine the latent trajectory bifurcations in the state space following presentation of stimuli and top-down cues in the network performing the one-modality DMS (i.e., one-modality RNN; Fig. 2e) and the two-modality DMS tasks (i.e., two-modality RNN; Fig. 2f–i). In the one-modality RNN, the trajectories during the first instruction period were separated into three pathways corresponding to the pro-DMS cue signal (light green and light purple in Fig. 2e), anti-DMS cue signal (dark green and dark purple in Fig. 2e), and the absence of the task cue instruction (i.e. late instruction condition; gray in Fig. 2e). The pro-DMS and anti-DMS pathways were then further bifurcated based on the identity of the first stimulus during the first stimulus period. For the two-modality RNN, we observed similar latent trajectories for the attended modality (Fig. 2f). However, the bifurcations driven by the identity of the first stimulus were not as well separated in the unattended modality (Fig. 2g). This pattern held true across pro- and anti-DMS trials, which exhibited qualitatively similar bifurcations (Fig. 2h,i). These findings were further confirmed by the cross-temporal discriminability analysis, which revealed that the identity of the first stimulus of the attended modality was more robustly encoded throughout the trial duration compared to the unattended modality (Fig. 2j–m; see *Methods*). Linear support vector machine (SVM) decoding analysis also revealed that the top-down cue signals were robustly encoded by the RNN models (Extended Data Fig. 1; see *Methods* for more details). These results suggest that lower-dimensionality representations of network dynamics sufficiently capture the network’s flexibility in encoding stimuli in a context-dependent manner, depending on stimulus relevance, as flexibly determined by top-down cues.

### Inhibitory units play a critical role in both encoding and maintaining top-down information

After establishing the model’s capacity for flexible encoding of top-down information, we next investigated the relative contributions of excitatory and inhibitory units in facilitating this process. Through dissection of the RNNs performing the one-modality and two-modality DMS tasks, we found that the optimized synaptic decay time constants for the excitatory units were significantly larger compared to the time constants of the inhibitory units (*p <* 0.001, two-sided Wilcoxon rank-sum test, Fig. 3a,b).

**Fig. 3.**
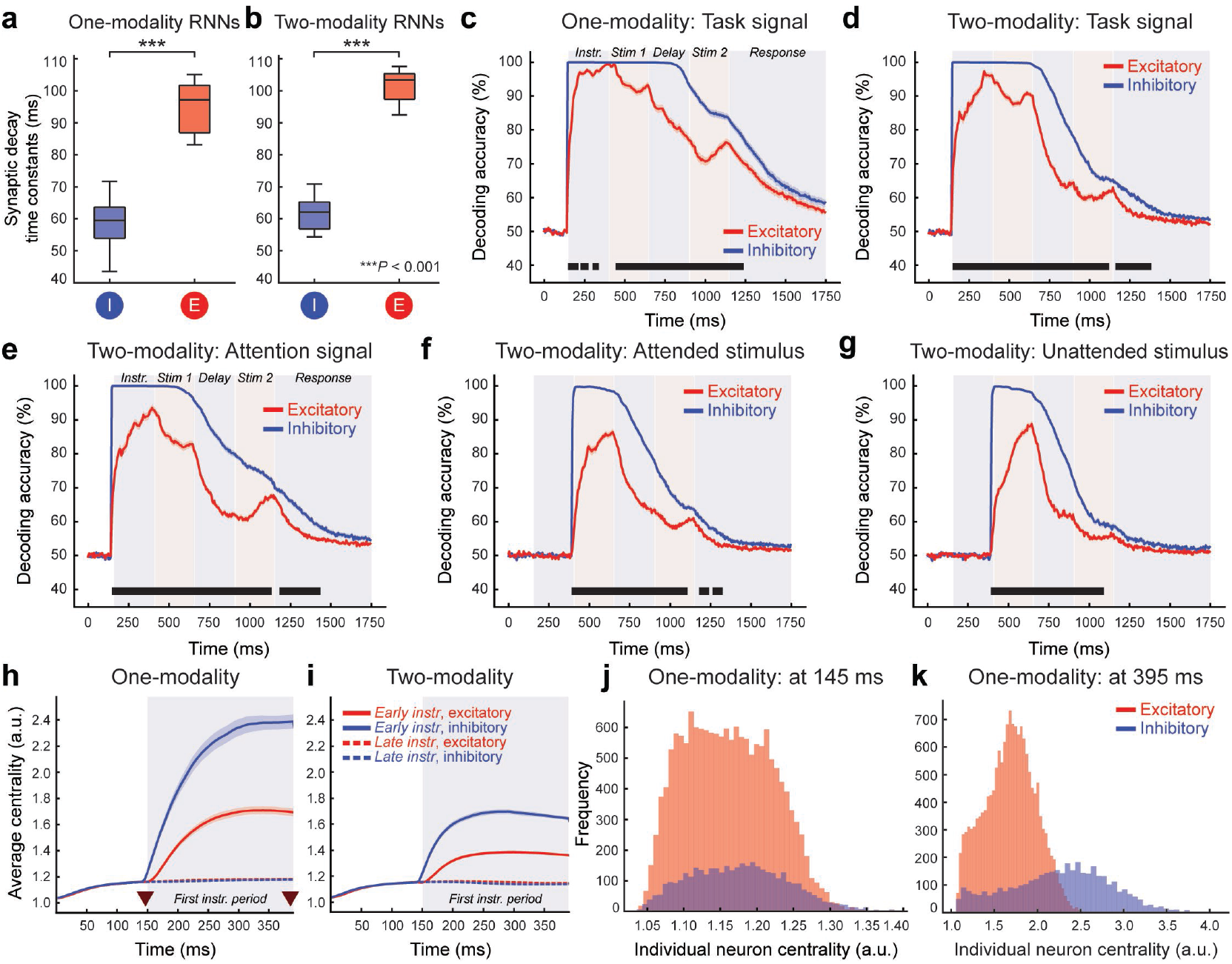
Inhibitory units robustly maintain top-down signals and establish functional connectivity hubs. **a-b.** Average synaptic decay time constants of inhibitory (blue) and excitatory (red) units in one-modality RNNs (**a**) and two-modality RNNs (**b**). **c.** SVM decodability of the task signal from inhibitory (blue) and excitatory (red) units from the one-modality RNNs. **d.** SVM decodability of the task signal from inhibitory (blue) and excitatory (red) units from the two-modality RNNs. **e.** SVM decodability of the attention signal from inhibitory (blue) and excitatory (red) units from the two-modality RNNs. **f.** SVM decodability of the attended first stimulus identity from inhibitory (blue) and excitatory (red) units from the two-modality RNNs. **g.** SVM decodability of the unattended first stimulus identity from inhibitory (blue) and excitatory (red) units from the two-modality RNNs. Black bars indicate significant differences between excitatory and inhibitory units (*p <* 0.05, two-sided KS test). **h.** Average functional connectivity centrality for all inhibitory (blue) and excitatory (red) units, across all one-modality RNNs, in early (solid) or late (dashed) instruction trials. Arrows indicate start and end of the first instruction period. Shaded areas indicate standard error of the mean. **i.** Same as **h**, for two-modality RNNs. **j.** Histogram of functional centrality values for all units in all one-modality networks at the beginning of the first instruction period. **k.** Same as **j**, at the end of the instruction period. Boxplot: central lines, median; bottom and top edges, lower and upper quartiles; whiskers, 1.5 × interquartile range; outliers are not plotted. \*\*\**p <* 0.001 by two-sided Wilcoxon rank-sum test.

Building on these findings, which indicate distinct roles for excitatory and inhibitory units, we conducted a series of linear support vector machine (SVM) decoding analyses. These analyses aimed to decode the task cue identity (anti- vs. pro-DMS) from the activity of excitatory and inhibitory units within RNNs engaged in the one-modality DMS task (see *Methods*). While the task cue identity could be readily decoded from the activities of both excitatory and inhibitory units following the cue presentation on early instruction trials, the SVM decoding accuracy was significantly higher for the inhibitory units throughout the trial duration (*p <* 0.05, 2-sample KS test, Bonferroni corrected across time; Fig. 3c). We observed similar trends for RNNs performing the two-modality DMS task (Fig. 3d). In these models, the inhibitory units also exhibited stronger maintenance of the attention signal compared to the excitatory units (Fig. 3e). Interestingly, decoding performance for stimulus identity was higher in inhibitory units for both the attended and unattended modalities, when compared to excitatory units (Fig. 3f,g).

Given these findings that highlight the distinct role of inhibitory units on maintaining task-related information, we next investigated their network-level implications through the functional connectivity patterns of inhibitory units. More specifically, we measured the functional closeness centrality of all units in each network across time (see *Methods*). Assuming that correlations between activation rates over time establish a metric of distance between units, it follows that units with greater centrality emerge as potential candidates for serving as functional hubs in their respective networks. While both excitatory and inhibitory units increased in centrality following the top-down cue presentation (*early instruction*) for the RNNs performing the one-modality and two-modality DMS tasks, the average centrality was higher for the inhibitory units after the onset of task and attention cue presentation (*p <* 0.05, two-sided 2-sample KS-test, Fig. 3h,i). Comparing the distribution of the centrality measure from individual units in the one-modality RNNs before (at 145 ms) and after (at 395 ms) the first instruction period on early instruction trials revealed that the centrality values of inhibitory units were distributed more widely post-instructions. This finding indicates the formation of inhibitory functional connectivity hubs following the top-down cue presentation (Fig. 3j,k). Taken together, these results suggest a central role for inhibitory units in encoding task-specific top-down information, as well as stimulus identity.

### Inhibitory-to-inhibitory connections carry top-down information

Given the crucial role of inhibitory units in encoding and sustaining top-down signals, we next investigated the neural circuitry surrounding these inhibitory units to further elucidate their contribution to the maintenance of task-specific top-down information.

Lesioning inhibitory connections to inhibitory units selective for the anti-DMS task cue (by reducing synaptic weights by 50%; see *Methods*) in a representative RNN trained to perform the one-modality DMS task resulted in the disruption of anti-DMS task signaling (Fig. 4a). Consequently, the network exhibited a tendency to perform the pro-DMS task irrespective of the task cue presented. By visualizing the neural dynamics of this lesioned RNN using CEBRA [63], we confirmed that the neural trajectories were no longer separated by the task cue condition during the first instruction period in early instruction trials (Fig. 4b).

**Fig. 4.**
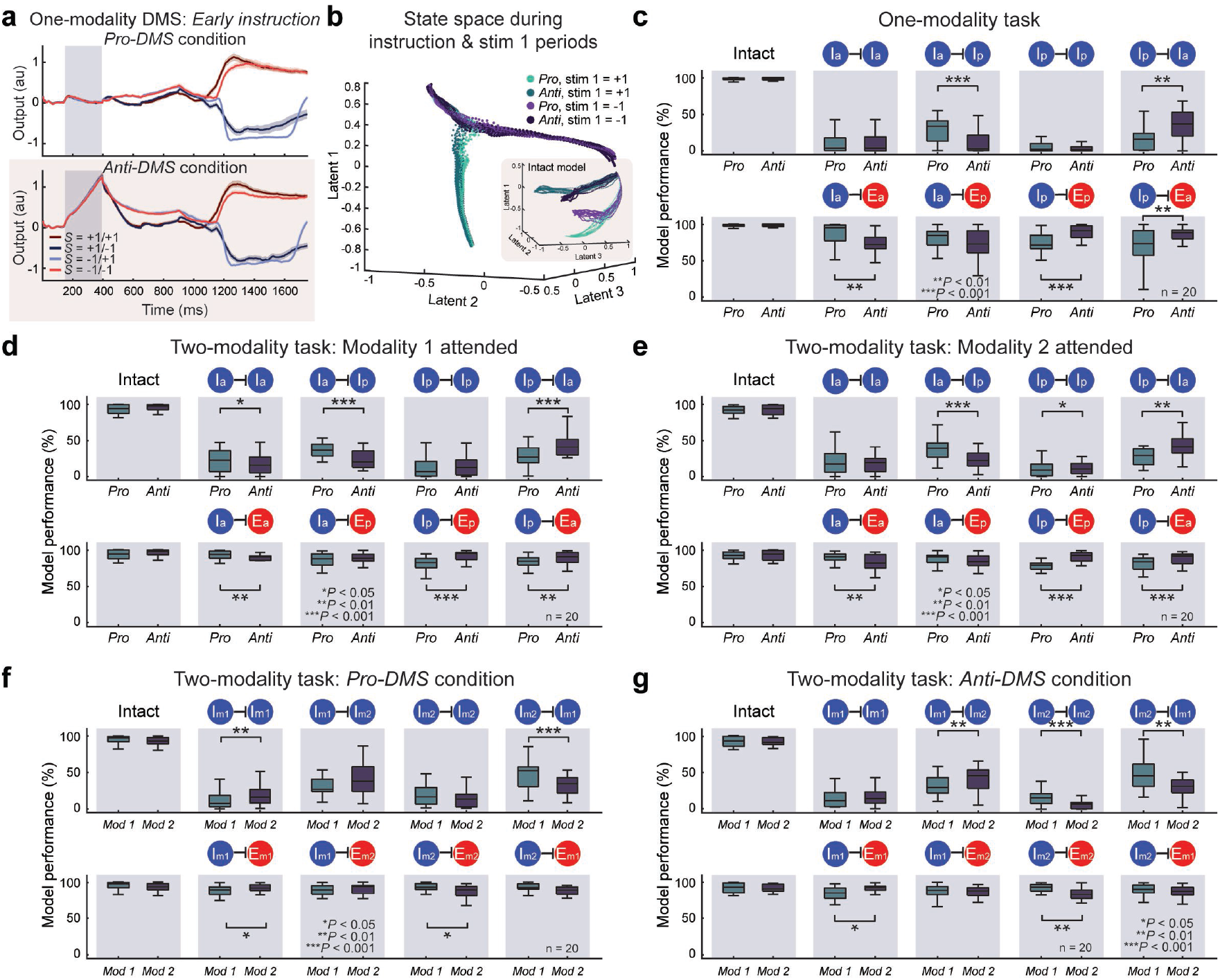
Inhibitory-to-inhibitory connections primarily encode task and attention cue signals. **a.** Reducing incoming inhibitory synaptic weights to inhibitory units selective to the anti-DMS cue by 50% in an example one-modality RNN led to the network performing the pro-DMS even when the anti-DMS cue was given (bottom). **b**) CEBRA neural trajectories of the lesioned model shown in **a** during the first instruction and first stimulus periods. Inset, trajectories from the intact network (identical to Fig. 2**e**). **c.** Performance of all 20 RNNs trained to perform the one-modality task when task-specific inhibitory-to-inhibitory (*I → I*; top) or inhibitory-to-excitatory (*I → E*; bottom) synaptic weights were reduced by 50%. **d.** Performance of all 20 RNNs trained to perform the two-modality DMS when task-specific *I → I* (top) or *I → E* (bottom) synaptic weights were reduced by 50% when modality 1 was attended. **e.** Same as **d**, when modality 2 was attended. **f.** Performance of all 20 RNNs trained to perform the two-modality DMS when modality-specific *I → I* (top) or *I → E* (bottom) synaptic weights were reduced by 50% when the task signal was fixed to pro-DMS. **g.** Same as **f**, when the task was fixed to anti-DMS. *I_a_*, inhibitory units preferring anti-DMS cue; *I_p_*, inhibitory units preferring pro-DMS cue; *E_a_*, excitatory units preferring anti-DMS cue; *E_p_*, excitatory units preferring pro-DMS cue; *I_m_*_1_, inhibitory units preferring modality-1 attention cue; *I_m_*_2_, inhibitory units preferring modality-2 attention cue; *E_m_*_1_, excitatory units preferring modality-1 attention cue; *E_m_*_2_, inhibitory units preferring modality-2 attention cue. Boxplot: central lines, median; bottom and top edges, lower and upper quartiles; whiskers, 1.5 × interquartile range; outliers are not plotted. \**p <* 0.05, \*\**p <* 0.01, \*\*\**p <* 0.001 by two-sided Wilcoxon rank-sum test.

Motivated by these findings, we employed a systematic approach where we lesioned synaptic connections based on task cue selectivity (pro- vs. anti-DMS) and unit type (excitatory vs. inhibitory) for the one-modality DMS task. We found that disrupting inhibitory-to-inhibitory (*I → I*) connections led to profound impairment in task performance (Fig. 4c top; \*\**p <* 0.01, \*\*\**p <* 0.001 by two-sided Wilcoxon rank-sum test). Disrupting similarly tuned inhibitory connections impaired performance in both pro- and anti-DMS trials (*Ia → Ia* and *Ip → Ip* in Fig. 4c top; \*\**p <* 0.01, \*\*\**p <* 0.001 by two-sided Wilcoxon rank-sum test). However, lesioning oppositely tuned *I → I* connections resulted in greater impairment in one type of task trials (*Ia → Ip* and *Ip → Ia* in Fig. 4c top; \*\**p <* 0.01, \*\*\**p <* 0.001 by two-sided Wilcoxon rank-sum test). We also observed significant changes in task performance when the *I → E* connections were lesioned (Fig. 4c bottom; \*\**p <* 0.01, \*\*\**p <* 0.001 by two-sided Wilcoxon rank-sum test). The effects of lesioning task cue-specific connections were similar in the RNN models trained to perform the two-modality task (Fig. 4d,e). In addition, *I → I* connections also carried attention cue-specific information in RNNs performing the two-modality DMS task (Fig. 4f,g).

These findings suggest that inhibitory-to-inhibitory connections are essential for integrating context-defining top-down information.

### Two-module RNN model with hierarchical organization

Recent empirical studies have shown that inhibitory-to-inhibitory connections within early sensory areas receive strong task-specific and context-based modulation by higher cortical areas [37, 54, 64].

Based on our results thus far, which underscore the significance of *I → I* connections in contextual coding, we next investigated whether inhibitory-to-inhibitory connections in low-level sensory areas would emerge as a critical target for task-oriented top-down modulation in our model. To accomplish this, we developed a two-module RNN model inspired by the cortical hierarchy. In this model, the first module (sensory module) is modeled after the early sensory cortex and receives the stimulus input signal only (Fig. 5a). The first module subsequently conveys the processed sensory signal to the second, non-sensory module through exclusively feedforward excitatory connections. Given previous findings from animal studies that showed lack of long-range feedforward inhibitory connections ([31]), we trained our two-module RNNs to perform the one-modality DMS task without intermodule feedforward inhibitory projections. The second module of the model, designed after higher cortical areas, receives the task cue signal (Fig. 5a). Using this architecture, we trained 20 two-module RNNs to perform the one-modality DMS task. On average, the two-module RNNs took longer to learn the one-modality task compared to the one-module networks (mean *±* stdev, 38,571 *±* 6,306 trials for the one-module RNNs vs. 55,221 *±* 10,187 trials for the two-module RNNs; Extended Data Fig. 2).

**Fig. 5.**
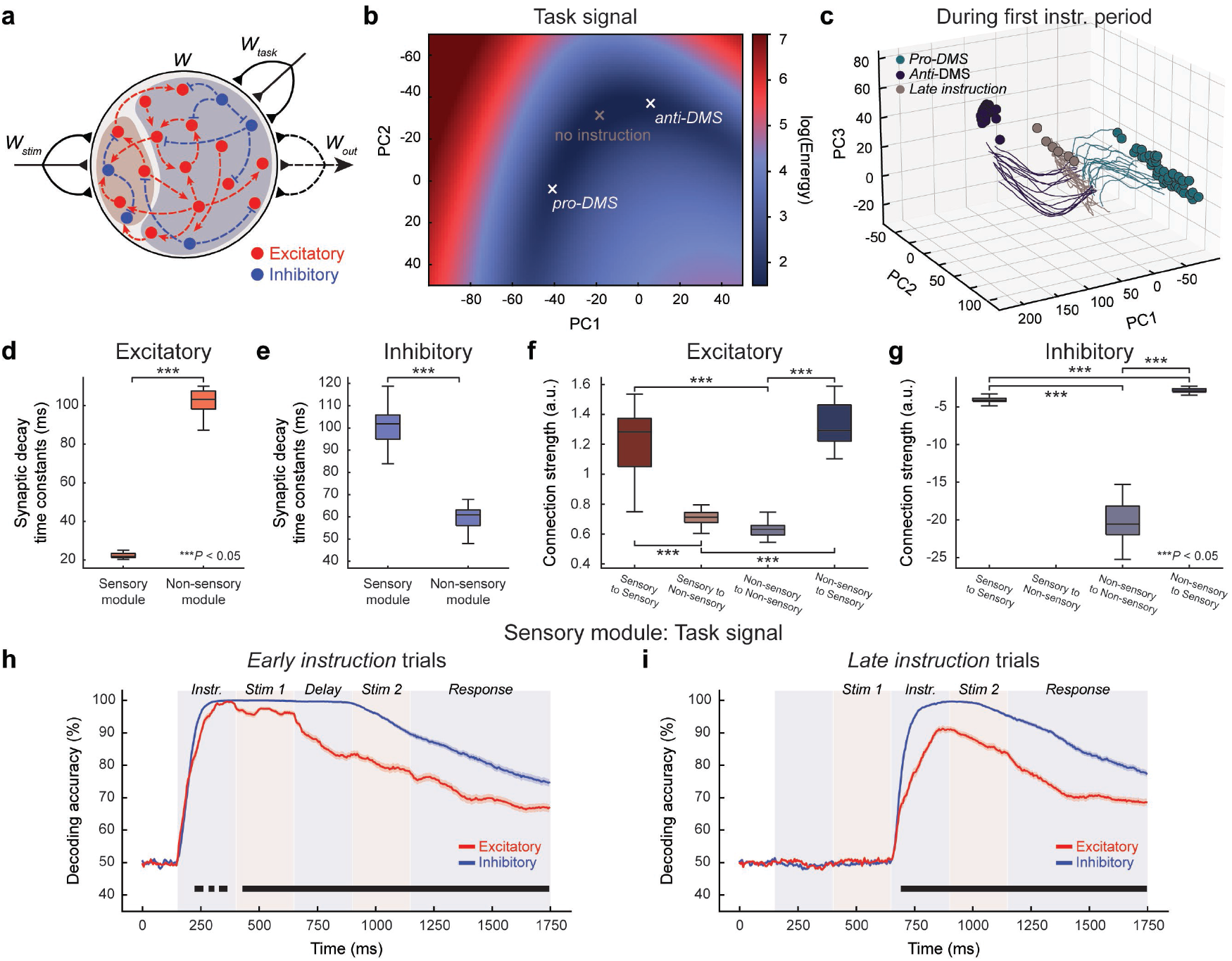
Dynamic coding and emergence of hierarchical inhibitory organization in two-module RNNs. **a.** Schematic of the two-module RNN model trained to perform the one-modality DMS task. The sensory module (red shade) contains 200 units and the non-sensory module (blue shade) contains 800 units. The sensory module receives the input stimulua signal, while the non-sensory module receives the task cue signal. There are no feedforward inhibitory projections from the sensory module to the non-sensory module. **b.** Heat map of kinetic energy in an example one-modality two-module RNN during the first instruction period. The middle cross indicates the local energy minimum for the late instruction condition (*no instruction*), while the left and right crosses indicate the new local energy minima for the pro- and anti-DMS conditions, respectively. **c.** Individual trial two-modality RNN trajectories in principal component space during the first instruction period. Principal component projections of fixed points are included. Trial task cues are divided into pro-DMS (teal), anti-DMS(dark blue) or no instructions (gray). **d-e.** Average synaptic decay time constants of excitatory units (**d**) and inhibitory units (**e**) in the sensory and non-sensory modules from 20 two-module RNN models trained to perform the one-modality DMS task. **f-g.** Comparison of average excitatory (**f**) and inhibitory (**g**) synaptic connection strengths within and between the sensory and non-sensory modules. **h.** Average decoding accuracy in early instruction trials for pro- vs. anti-DMS signal in the sensory module of two-module networks, including either excitatory (red) or inhibitory (blue) units. Shaded areas indicate standard error of the mean. Black bars indicate significant differences between excitatory and inhibitory units (*p <* 0.05, two-sided KS test). **i.** Same, for late instruction trials. Boxplot: central lines, median; bottom and top edges, lower and upper quartiles; whiskers, 1.5 × interquartile range; outliers are not plotted. \*\*\**p <* 0.001 by two-sided Wilcoxon rank-sum test (**d** and **e**) or Kruskal-Wallis test followed by Dunn’s multiple comparison test (**f** and **g**).

To investigate if the two-module RNNs processed the task cue signal in a manner similar to that observed in the one-module RNNs, we characterized the kinetic energy landscape and performed the fixed-point analysis on the two-module networks during the first instruction period (150-400 ms; see *Methods*). Similar to the landscape and energy minima seen in the one-module RNNs (Fig. 2a), we observed an inverted U-shaped energy landscape and displacement of energy minima as a function of the task cue identity (Fig. 5b). In addition, the fixed point analysis revealed a line attractor structure which was also displaced based on the task cue signal presented to the network (Fig. 5c). These results suggest that network trajectories are conditioned by a systematic shift in fixed point structure, which is brought about by the presence of distinct top-down signals in the form of task instructions.

Characterizing the synaptic decay dynamics of the two-module RNNs revealed slower excitatory dynamics in the non-sensory module compared to the sensory module (Fig. 5d; *p <* 0.001 by two-sided Wilcoxon rank-sum test). The inhibitory units exhibited the opposite trend (Fig. 5e; *p <* 0.001, two-sided Wilcoxon rank-sum test). Investigating the average synaptic connection strength revealed strong excitatory connections within the sensory module and from non-sensory to sensory modules (Fig. 5f; *p <* 0.001, Kruskal-Wallis test followed by Dunn’s multiple comparison test). Interestingly, the average inhibitory signaling was the strongest within the non-sensory module (Fig. 5g; *p <* 0.001, Kruskal-Wallis test followed by Dunn’s multiple comparison test). Both *I → E* and *I → I* connections within the non-sensory module were significantly stronger than their counterparts in the sensory module (Extended Data Fig. 3). These findings are consistent with previous empirical evidence suggesting increased inhibitory strength and diversity along the cortical hierarchy [53, 65].

Linear SVM decoding analyses in the two-module RNNs showed that the task signals were more readily decodable from the inhibitory units than from the excitatory units in both sensory and non-sensory modules (Fig. 5h,i; see Extended Data Fig. 4 for the non-sensory module results, *p <* 0.05, two-sided KS test with Bonferroni corrected across time). Most notably, the SVM decodability of the task signal decayed slower for the inhibitory units compared to the excitatory units in both modules.

### Inhibitory-to-inhibitory connections in both sensory and non-sensory modules encode top-down information

Analyzing the neural trajectories of an example two-module RNN during the first stimulus period for early instruction trials revealed trajectory grouping based on the task cue identity (Fig. 6a). Lesioning *I → I* connections in both sensory and non-sensory modules by reducing the weights by 50% abolished this top-down modulation (Fig. 6b), mirroring the results observed in the one-module RNN (Fig. 4b). Lesioning other synaptic connections (i.e., *E → E, E → I, I → E*) did not lead to disruption of the top-down information encoding. Thus, these results suggest that the hierarchical, two-module RNNs also rely on inhibitory-to-inhibitory connections to encode contextual information.

**Fig. 6.**
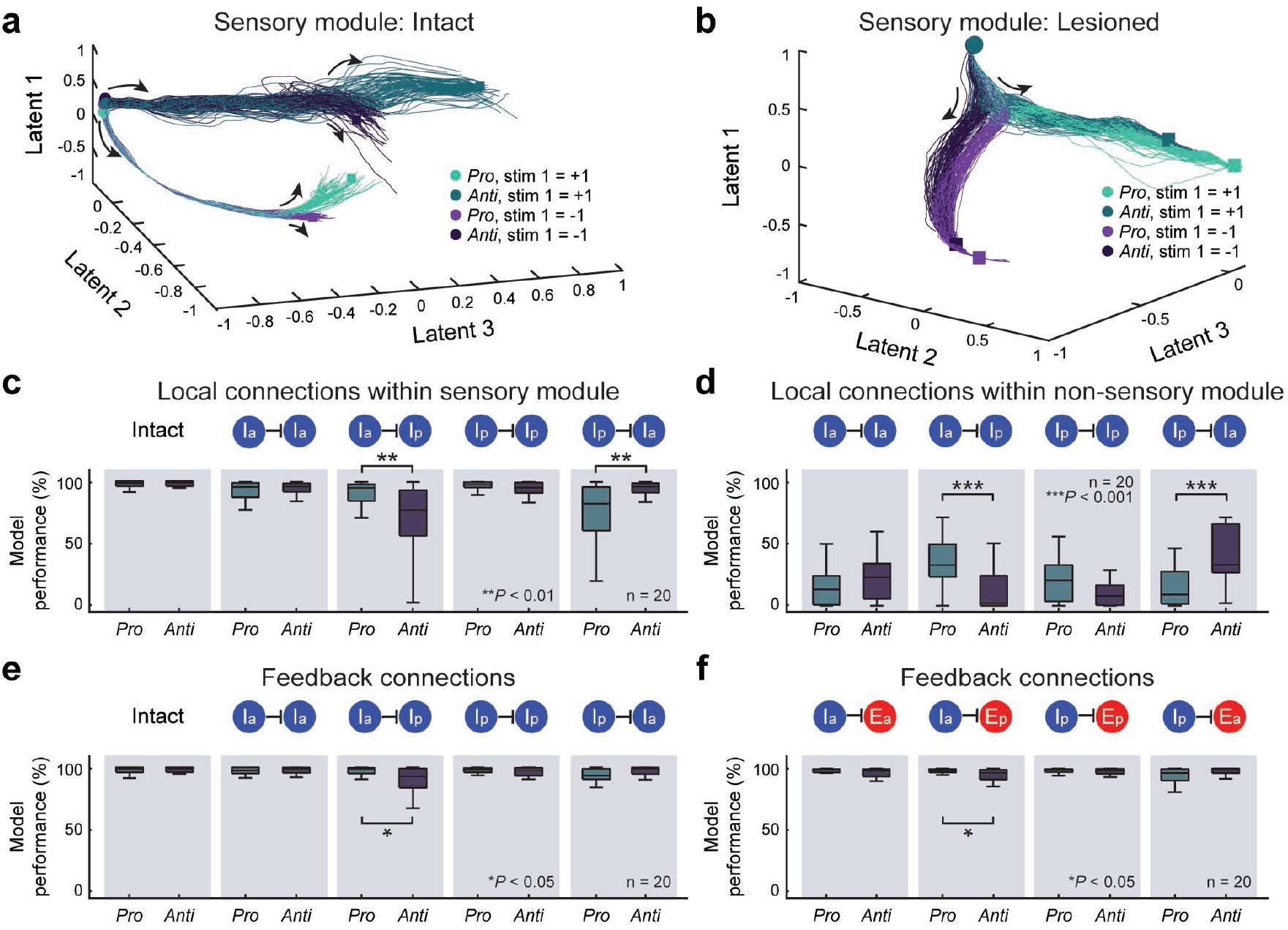
*I → I* connections in both sensory and non-sensory modules contribute to task cue encoding. **a.** CEBRA neural trajectories during the first instruction and first stimulus periods from an example two-module RNN. **b.** CEBRA trajectories during the first instruction and first stimulus periods from the same RNN model shown in **a**, but with *I → I* connections lesioned. **c-d.** Performance of all 20 two-module RNNs trained to perform the one-modality DMS task when task-specific *I → I* connection strengths within the sensory module (**a**) or within the non-sensory module (**b**) were reduced by 50%. **e-f.** Performance of all 20 two-module RNNs trained to perform the one-modality task when task-specific *I → I* connection strengths (**c**) or *I → E* (**d**) from the non-sensory to the sensory module were reduced by 50%. Boxplot: central lines, median; bottom and top edges, lower and upper quartiles; whiskers, 1.5 × interquartile range; outliers are not plotted. \**p <* 0.05, \*\*\**p <* 0.001 by two-sided Wilcoxon rank-sum test.

Performing a systematic lesioning analysis, similar to the one utilized for the one-module RNNs in Figure 4, demonstrated the importance of inhibitory units with opposite tuning to the task cue (i.e., anti- vs. pro-DMS) in the sensory module for reliably encoding the task signal (Fig. 6c). In addition, disrupting *I → I* connections within the non-sensory module severely impaired the task performance of all the RNNs, confirming that top-down information is primarily encoded by *I → I* connections in the non-sensory module (Fig. 6d). Lesioning feedback inhibitory connections had minimal influence on the task performance (Fig. 6e,f).

These results suggest that the task cue information is encoded by *I → I* synapses in the non-sensory module. What is even more unexpected is that the top-down information is transmitted to the sensory module via feedback connections and encoded by *I → I* connections within the sensory module (Fig. 6a). Although some connections did not reach statistical significance, based on the observed trend, it is likely that *I → E* connections play a significant role in transmitting top-down information from the non-sensory module to the sensory module (Fig. 6f).

### Disinhibitory circuits in mouse primary visual cortex and RNN sensory module encode top-down information

Building on the findings from the previous section, which indicate the role of sensory areas in encoding both stimulus-driven (bottom-up) and task-related (top-down) information, we proceeded to test our hypothesis. Specifically, we proposed that cortical *I → I* connections within early sensory processing regions represent top-down information via feedback from higher cortical areas. To investigate this, we analyzed a publicly available experimental data which demonstrated the capability of the mouse primary visual cortex (V1) in representing contextual information [54]. This electrophysiology dataset was collected with a linear array recording electrode covering all layers of V1 and preprocessed with standard filtering and spike sorting pipelines.

In this study, during V1 activity recording, mice were trained to identify the location of a salient grating visual pattern (“figure”) that stood out from the visual background (“ground”) [54]. These patterns were distinguished based on either orientation, phase, or texture (Fig. 7a). Crucially, in each of these conditions, the sensory inputs chosen to fall within the exact receptive fields of the recorded neurons remained strictly identical in both the figure and background conditions (see *Methods*). This implies that any observed difference in figure versus ground coding within V1 neurons representing these receptive fields would stem from coding dependent on top-down feedback. This mirrors our approach in configuring our RNNs to process identical bottom-up sensory information under varying contexts informed by top-down cues. Therefore, any disparity in neural activity observed across conditions in the mouse dataset would be attributed to the context indicated by stimulus regions outside of the measured neural receptive field.

**Fig. 7.**
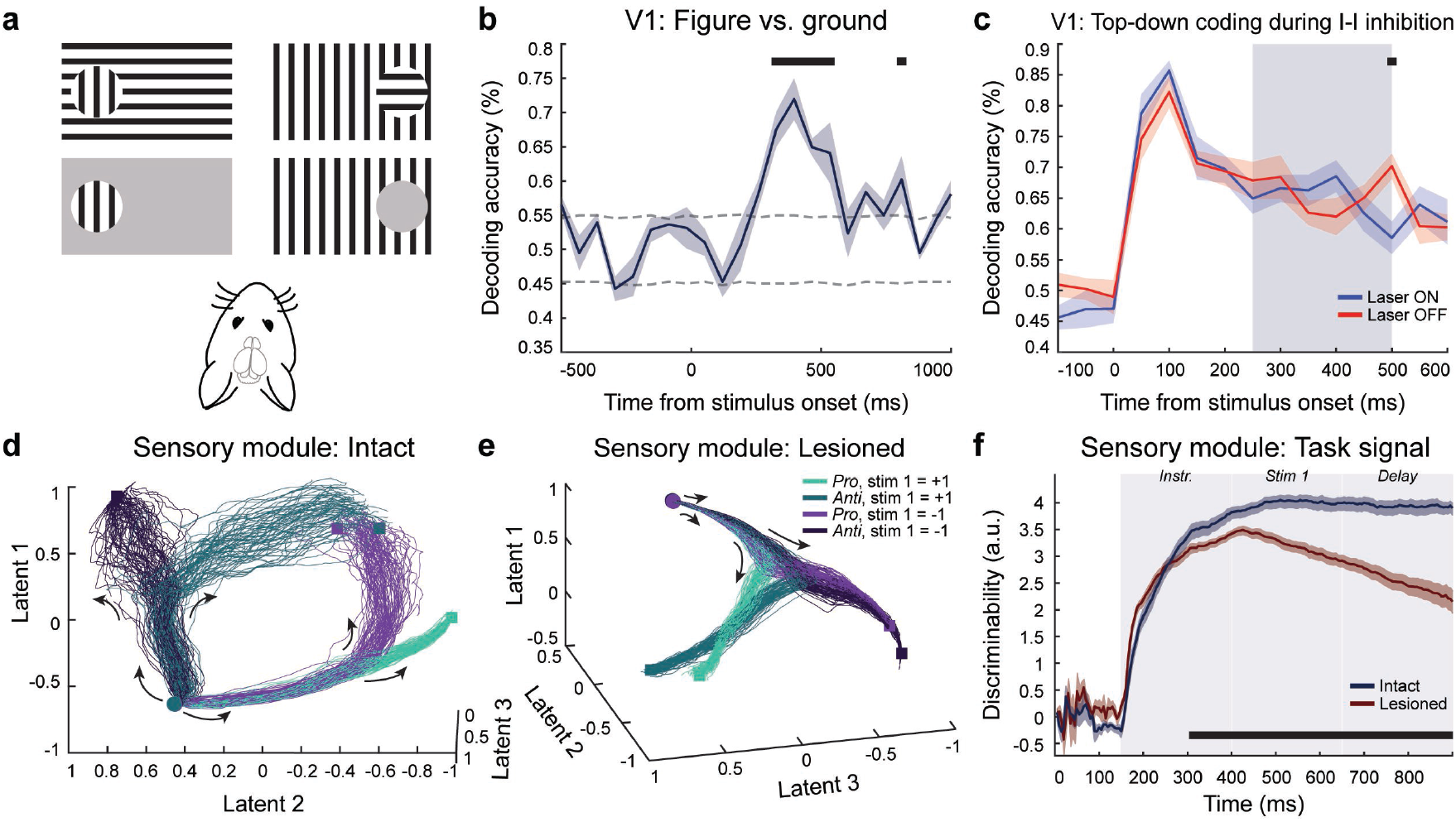
Top-down encoding in mouse V1 and RNN sensory module. **a.** Task summary. In the orientation condition, mice viewed an orientation pattern as either part of a figure (top left) or as part of the background (top right). Phase and texture detection conditions (not shown) were also included in the analysis. In control trials (bottom row), mice must contrast a target against a gray background which did not require figure-ground segregation. This design ensured that the same sensory information within receptive fields could possess different contextual meanings (figure vs. ground). Reproduced from [54]. **b.** SVM decoding of top-down figure vs. ground signal from V1 neurons, holding sensory input constant. Black bars indicate significant decoding (*p <* 0.05, random permutation test). Dashed lines indicate 95% confidence intervals from permutations. **c.** SVM decoding of top-down signal during optogenetic I-I inhibition in V1, in laser on (blue) and laser off (red) trials. Gray bar indicates laser on time window. Black bar indicates significant difference in decoding across conditions (*p <* 0.05, one-sided KS test). **d.** CEBRA trajectories within the sensory module of an example two-module RNN during the first stimulus period when the task signal was presented early. **e**. Same as **d** but when the connections from the non-sensory module to the sensory module were completely removed. **f.** Comparison of average within-time temporal discriminability for the first stimulus identity across 20 trained, two-module RNNs under two conditions: one with intact connections including feedback and the other without feedback (lacking connections from non-sensory to sensory modules).

First, we tested whether the activity of V1 neurons could differentiate between figure and ground conditions, even when the sensory information within their receptive fields remained identical. A successful figure vs. ground decodability would be an evidence for feedback from regions outside receptive fields into the measured V1 sites. Performing a linear SVM decoding analysis on the V1 activity to decode the condition label (figure vs. ground) revealed that the V1 responses were modulated by the task context (Fig. 7b; *p <* 0.05, permutation test, Bonferroni corrected). The high decoding accuracy indicates that feedback signals from regions outside the receptive fields robustly modulate V1 activity in response to information about task context.

In sensory cortical areas, including V1, vasoactive intestinal peptide-expressing (VIP) interneurons send inhibitory projections to somatostatin-positive (SST) interneurons. This forms micro-circuitry characterized by inhibitory-to-inhibitory connections, similar to the *I → I* connections observed in the sensory module of our two-module RNN model (Fig. 5a). Leveraging the experiments within the dataset where VIP interneurons of mouse V1 were optogenetically silenced, we analyzed the role of VIP neurons in figure-background discrimination. Through the SVM decoding analysis, we observed a decrease in decoding accuracy following optogenetic silencing of V1 VIP neurons (*p <* 0.05, one-sided Kolmogorov-Smirnov test, Bonferroni corrected; Fig. 7c). These results demonstrate the importance of intact inhibitory-inhibitory projections for successfully encoding top-down information in V1 circuits.

We next utilized CEBRA [63] to analyze the neural dynamics of the sensory module in an example two-module RNN model during the first stimulus presentation in early instruction trials of the one-modality DMS task. The recovered latent space revealed distinct neural trajectories delineated by the task cue signal (Fig. 7d). The trajectories were initially divided into two arms based on the identity of the task cue (light green and light purple vs. dark green and dark purple trajectories in Fig. 7d). Within each arm, the trajectories were further separable by the first stimulus identity (+1 or -1). Removing all the inhibitory-to-inhibitory connections within the sensory module abolished the effects of the task signal in the sensory module (Fig. 7e). These results highlight a convergence between theoretical models and experimental evidence for the importance of feedback into lower sensory areas in coding top-down information.

Performing the cross-temporal discriminability analysis on the sensory module across all 20 trained two-module RNNs demonstrated that the task cue identity could be reliably decoded, providing additional confirmation of the top-down modulation imposed by the non-sensory module (Fig. 7f; see *Methods*). When the sensory *I → I* connections were removed, the task cue identity could not be decoded as reliably as in the intact model (Fig. 7f), indicating the crucial role of these connections in facilitating top-down modulation within the network architecture.

### Introducing retro-cue condition elicits shift in strategy

Identifying relevant sensory stimuli and determining the timing of their relevance is often dynamic in the real world. To better capture this, we modified the two-modality DMS task by introducing a retro-cue condition as illustrated in Figure 8a. The task still involves two streams of stimuli (modality 1 and modality 2) along with the task cue (pro- and anti-DMS). In this task version, the task cue was fixed to be given before the first stimulus; see *Methods*). Notably, in this modified version, the attention cue may be presented before the first stimulus (*early instruction*), after the first stimulus (*late instruction*), or after the second stimulus (*retro-cue*), as depicted in Figure 8a. This modification provides a more nuanced exploration of attentional processes by allowing flexibility in the timing of attention cue presentation.

**Fig. 8.**
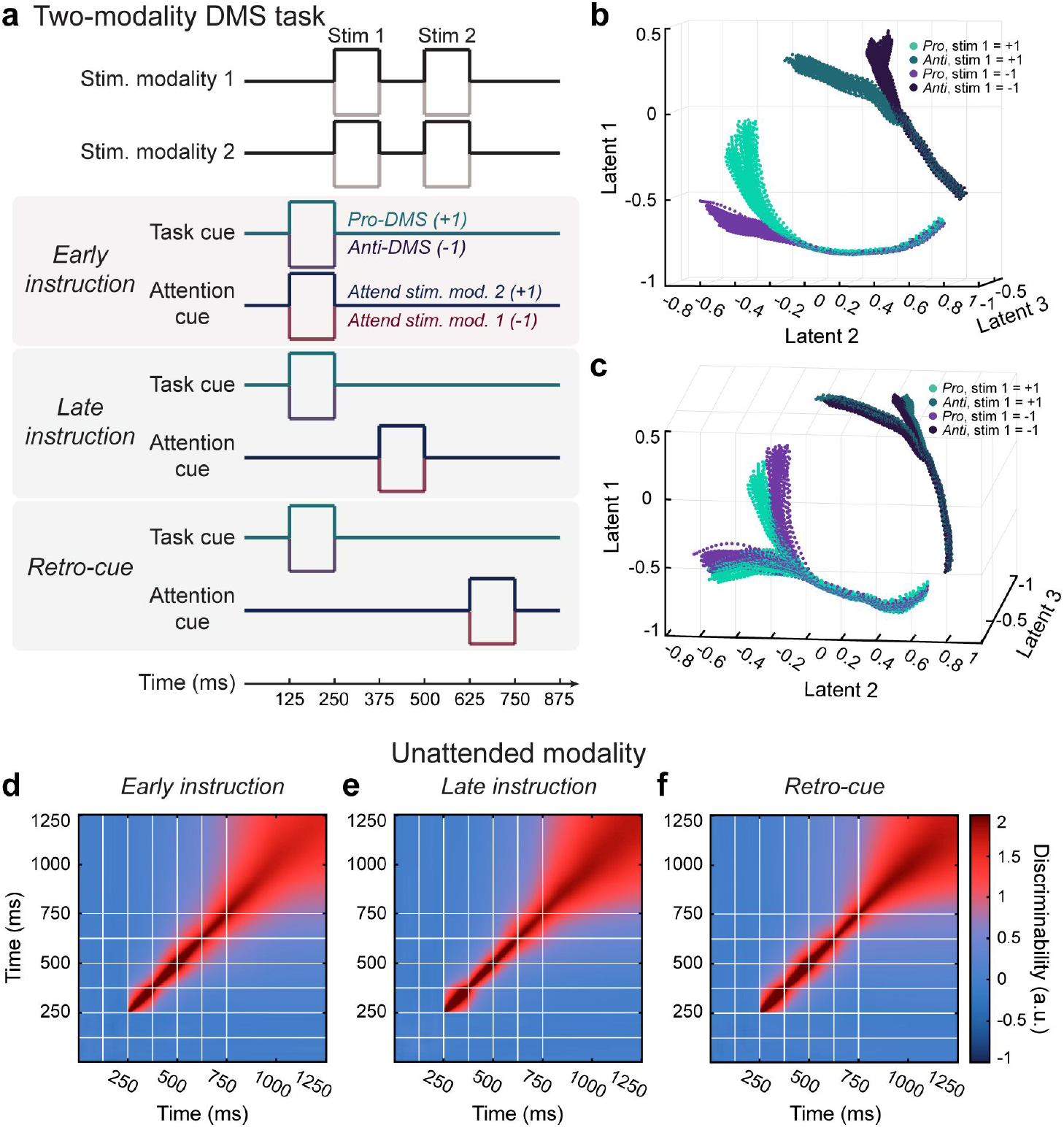
Retro-cue signal forces the model to maintain both unattended and attended sensory signals. **a.** Schematic of a modified version of the two-modality DMS task which includes a retro attention cue signal. The task cue signal was always given early (before the onset of the first stimulus), while the attention cue was given before the first stimulus (*early instruction*), during delay (*late instruction*), or after second stimulus (*retro-cue*). **b.** CEBRA trajectories during the first instruction and first stimulus periods for the attended modality when the attention cue signal was given early (example RNN). **c.** Same as **b** but for the unattended modality. **d-f.** Average temporal discriminability of the first stimulus identity of the unattended modality when the attention cue was given early (**d**), late (**e**), or after the second stimulus (retro; **f**).

We trained 20 one-module RNNs to perform this modified two-modality DMS task. We then compared the resulting network dynamics for the case where top-down cues (both task and attention cue signals) were given before the first stimulus against the dynamics observed from the RNNs trained for the original two-modality DMS task (Fig. 2). Applying CEBRA to an example RNN trained for the modified two-modality DMS task revealed that the trajectories for the attended modality closely resembled those obtained from the example RNN trained for the original two-modality DMS (i.e., compare Fig. 8b with Fig. 2f). The trajectories were initially distinguished by the task cue (pro- vs. anti-DMS). Subsequently, each task “arm” bifurcated based on the identity of the attended first stimulus (Fig. 8b). Interestingly, in the case of pro-DMS, where the unattended first stimulus had an identity of +1, the trajectories diverged based on the identity of the attended first stimulus (illustrated by the green trajectories in Fig. 8c). This stands in stark contrast to the trajectories observed in the RNN model performing the original two-modality DMS task, where the trajectories of the unattended modality were unaffected by the attended modality (Fig. 2g). In other words, the introduction of the retro-cue condition to the two-modality task forced the RNNs to maintain not only the attended but also the unattended stimuli throughout the trial period. This is further underscored by the robust temporal discriminability of the first stimulus of the unattended modality across the three attention cue conditions (i.e., early, late, and retro cue conditions) in the 20 RNNs trained for the modified two-modality DMS task (Fig. 8d–f; compare these to Fig. 2l,m).

Overall, introducing the retro-cue condition resulted in the RNN model employing a different strategy to flexibly prioritize and maintain relevant information throughout the task execution.

## Discussion

The present study provides a unique set of cognitive tasks that require flexible stimulus-response mapping and enable investigation into the interaction between bottom-up and top-down signaling. By constructing and training biologically constrained recurrent neural networks (RNNs) on these tasks, we elucidate and propose possible neural mechanisms essential for such flexible information processing and cognitive control. Importantly, we show that selective inhibition plays a central role in shaping network dynamics during context-dependent decision-making. Moreover, through the utilization of hierarchically structured RNN models (i.e., two-module RNNs) and data from the mouse primary visual cortex, we demonstrate the involvement of selective inhibition in encoding top-down information within early sensory areas. Together, these findings offer a complete computational and empirical account for the role of top-down feedback into lower level sensory areas.

Previous experimental and computational studies have highlighted the significance of disinhibitory circuits governed by inhibitory-to-inhibitory signaling in performing high-level cognitive functions [33, 40, 66–68]. For instance, disinhibitory effects exerted by vasoactive intestinal peptide–expressing (VIP) and somatostatin-positive interneurons (SST) in higher-level cortical areas, such as the prefrontal cortex, have been shown to be critical for working memory and social memory [52, 69]. While recent animal studies have started to explore how these mechanisms are utilized for other cognitive functions [70–72], the role of inhibitory-to-inhibitory connections in facilitating flexible cognitive switching remains poorly understood. This ambiguity is primarily attributed to the lack of computational models incorporating relevant biological constraints, coupled with scarcity of cognitive tasks that allow for orthogonal manipulation of sensory and temporally varying top-down signals. To bridge this gap, we trained RNNs composed of model neurons to perform cognitive tasks requiring adaptive switching. Through several computational analyses, including lesion studies, we demonstrate that a subset of *I → I* synapses naturally emerge in our model to adeptly map sensory information into context-dependent decisions. Consistent with our findings, recent animal studies have identified disinhibitory circuit motifs facilitating flexible routing of sensory information [33, 37, 38, 54, 72].

Particularly noteworthy are our findings that *I → I* connections within the sensory module of our two-module RNN model encode top-down information, and that these connections are necessary for context-dependent mapping of the sensory information. Furthermore, we reported that feedback signals from higher-level cortical areas contribute to emergence of these *I → I* connections in early sensory areas. These findings indicate a hierarchical organization of inhibitory circuits involved in encoding and processing top-down information. In support of this, Kirchberger et al. demonstrated the critical role of feedback inputs from higher visual areas in generating context-dependent signals in the mouse primary visual cortex through optogenetic manipulation [54]. While the authors observed reduced context-encoding upon optogenetic silencing of excitatory neurons in the higher visual cortex [54], the specific circuitry underlying these feedback projections warrants further investigation.

The possibility that aspects of complex computations, typically ascribed to higher cortical areas, may also occur in lower-level areas challenges the prevailing notion of the cognitive limitations of lower-level areas. Building in representational redundancies throughout cortical hierarchies allows for complex computations to occur in both lower sensory areas and higher-level areas [25, 41, 42]. This mechanism enhances the robustness of representations and facilitates efficient processing, enabled by top-down feedback signals guiding multiplexing in sensory areas.

Building on the insights gained from this study, future directions could explore whether similar circuit mechanisms are employed across various cognitive domains, potentially uncovering universal circuit motifs underlying flexible sensory processing and cognitive control across various contexts. Furthermore, enhancing the biological realism of computational models and RNNs by incorporating spiking model neurons and biologically plausible learning rules could unveil other neural mechanisms based on precise timing-based computations. Finally, validating our experimentally testable hypotheses and predictions using neural data from both non-human primates and humans, particularly epilepsy patients with depth electrodes implanted for seizure monitoring and potential intervention, represents a crucial future direction. This step would provide an improved understanding of representational redundancies within cortical networks and help identify compensatory mechanisms that preserve flexible computation in networks disrupted by disorders. Additionally, these insights will inform the development of targeted interventions and therapeutic strategies for neurological disorders that impair sensory processing and cognitive control.

## Acknowledgements

We thank Jorge Aldana and the Computational Neurobiology Laboratory at the Salk Institute for assistance with computing resources. We thank Lisa Kirchberger of the Netherlands Institute for Neuroscience for the support in providing access to mouse data for this manuscript. We are also grateful to Xiao-Jing Wang and Mackenzie Mathis for their invaluable feedback on our work. This work was funded by the ARL Human Guided Intelligent Systems grant (W911NF-23-2-0067) and the Strengthening Teamwork for Robust Operations in Novel Groups (STRONG) grant (W911NF-22-2-0148). The funders had no role in study design, data collection and analysis, decision to publish, or preparation of the manuscript.

## Author contributions

R.K. and N.R. conceived and designed the research; T.G.A., R.K., and N.R. analyzed data; T.G.A., R.K., and N.R. wrote the manuscript.

## Methods

### Continuous-rate recurrent neural network (RNN) model

In this study, we constructed the following continuous-variable recurrent neural network (RNN) model:

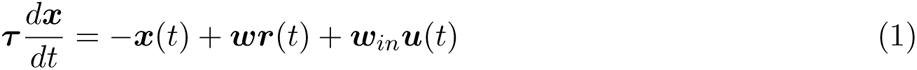

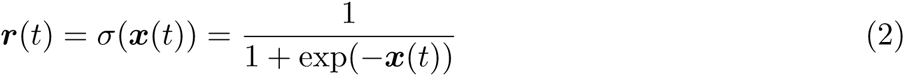

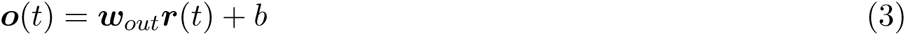

where ***τ*** *∈* ℝ^1^*^×N^* refers to the synaptic time constants, ***x*** *∈* ℝ*^N×T^* denotes the synaptic current variable from from *N* units across *T* time points. By applying a sigmoid nonlinearity, we estimated the firing rates ***r*** *∈* ℝ*^N×T^* based on the synaptic current values (***x***). The connection weights from the time-varying inputs (***u*** *∈* ℝ*^Nin×T^*; *N_in_*, the number of input channels) to the network were represented by the weight matrix ***w****_in_ ∈* ℝ*^N×Nin^*. Additionally, ***w*** *∈* ℝ*^N×N^* contains connection weights between the *N* units.

To compute the network output (***o*** *∈* ℝ^1^*^×T^*), we linearly combined all the firing rates specified by the output connection weight matrix, ***w****_out_ ∈* ℝ^1^*^×N^*, along with the constant bias term, *b*. The network size (*N*) was set to 1000 for all the networks.

For the one-modality DMS task, the input signals (***u***) contained three channels: two channels for the two sequential stimuli and one channel for the task cue signal (Fig. 1a). For the two-modality DMS task, ***u*** contained 6 channels, corresponding to two stimulus modalities (two channels for each modality), one channel for the task cue signal, and another one for the attention cue (Fig. 1b). For the one-module RNNs, ***u*** was projected to all the units in the network. However, for the two-module RNNs, the stimulus signals were injected into the first 200 units, while the task and attention cues were provided to the second module comprising 800 units (Fig. 5a).

For the two-module RNN model, the recurrent connectivity weight matrix (***w***) was constrained to remove the feedforward inhibitory connections from the first module (first 200 units; sensory module) to the second module (non-sensory module) by applying a binary mask.

### Training and task details

The dynamics in Eq. (1) were discretized using the first-order Euler approximation method and using the step size (Δ*t*) of 5 ms:

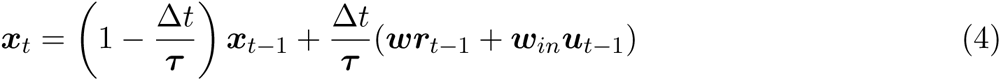

where ***x****_t_* = ***x***(*t*). Given that the units in this network model communicate through differentiable and continuous signals, a gradient-descent supervised method, known as backpropagation through time (BPTT; [73]), was employed to train our RNNs to perform cognitive tasks. Specifically, the trainable parameters of the model included ***w***, ***τ***, ***w****_out_*, and *b*. We used Adam (adaptive moment estimation) optimization algorithm to update these parameters. The learning rate was set to 0.01, and the TensorFlow default values were used for the rest of the parameters including the first and second moment decay rates. To further impose biological constraints, we enforced Dale’s law (uniform neurotransmitter release characteristics within separate excitatory and inhibitory neurons) using methods similar to those implemented in previous studies ([43, 49, 74]). To adhere to empirical findings regarding the ratio of excitatory and inhibitory neurons observed in the brain, each RNN consists of 80% excitatory and 20% inhibitory units (i.e., E-I ratio of 80/20; [30, 75, 76]).

Importantly, instead of fixing the synaptic decay constant (***τ***) to a fixed value for all the units, we optimized the parameter for each unit. The parameter was trained to range from 20 ms to 125 ms to model heterogeneous synaptic dynamics of different receptors in the cortex [77, 78]. We initialized the synaptic decay time constant parameter (***τ***) using

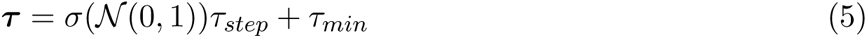

where *σ*(*·*) is the sigmoid function and *N* (0, 1) refers to the standard normal Gaussian distribution. *τ_min_* = 20 ms and *τ_step_* = 105 ms were used to constrain the parameter to range from 20 ms to 125 ms.

Our model training was deemed successful if the following two criteria were satisfied within the first 70,000 epochs: 1) loss value (defined as the root mean squared error between the network output and target signals) *<* 7; and 2) task performance *>* 95% for the one-modality DMS task and *>* 85% for the two-modality DMS task. If the network did not meet the criteria within the first 70,000 epochs, the training was terminated. For each successfully trained RNN, we simulated 400 trials for the one-modality task and 1,000 trials for the two-modality task, randomizing each trial to contain one possible combination of instruction order, top-down cues, and stimulus identities. Using these criteria, we trained 20 RNNs for each task.

#### One-modality DMS task

For the one-modality DMS task, the trial duration was set to 350 steps with 5 ms step size, and the input matrix (***u*** *∈* R^3^*^×^*^350^) contained two input channels for the two sequential stimuli (Stim 1 and Stim 2 in Fig. 1a). The third channel was used to deliver the task cue signal. The first channel was employed to present Stim 1 (lasting 250 ms) at 400 ms, and the second channel delivered Stim 2 (lasting 250 ms) 250 ms after the offset of the first stimulus. During the stimulus period, the input channel was set to either -1 or +1. For the early instruction condition, the task cue channel was set to either -1 or +1 from 150 ms to 400 ms. For the late instruction condition, the task cue channel was adjusted to -1 or +1 during the delay period, spanning from 650 ms to 900 ms.

#### Two-modality DMS task

The input matrix for the two-modality task contained 6 input channels (***u*** *∈* ℝ^6^*^×^*^350^). The initial two channels were employed to present two sequential stimuli for the first modality, whereas the subsequent two channels delivered stimuli for the second modality. The fifth channel was dedicated for the task cue signal, while the last channel was used to present the attention cue. For the early instruction condition, both task and attention signals were delivered at 150 ms. For the late instruction condition, the task and attention signals were presented at 650 ms.

#### Modified two-modality DMS task

For the modified version of the two-modality DMS task (Fig. 8), the input matrix closely resembled that of the standard two-modality task (see above). The trial duration for this variant was reduced to 250 steps (1,250 ms), and the stimulus window was also shortened to 125 ms. In this task, the task cue signal was always delivered before the first stimulus at 125 ms. However, the attention cue signal was randomly delivered either before the first stimulus (*early instruction*), after the first stimulus at 375 ms (*late instruction*), or after the second stimulus at 625 ms (*retro-cue*).

### SVM decoding and state space analyses

We performed decoding on RNN activity with a 5-fold cross-validated linear support vector machine analysis (SVM; *sklearn* function SVC). Given a trained RNN model, we first generated firing rate timecourses (***r*** in Eq. (3)) of 1,000 units across 1,000 simulated trials. Next, the data were randomly sampled for instruction order (early vs. late), stimulus identity (+1 vs. -1), and top-down task signals (pro-DMS vs. anti-DMS; as well as attended vs. unattended for the RNNs performing two-modality DMS). The SVM classifier was trained on the firing activities from either excitatory or inhibitory units and then tested on the left-out trials to distinguish each of the signals of interest.

To characterize the network dynamics in response to distinct external inputs, we performed a fixed-point analysis [60] using the FixedPointFinder toolbox [61]. This method involves numerically minimizing a proxy energy function derived from the RNN update equations to identify fixed points and subsequently obtaining linearized dynamics around these points to describe their behavior. In our case, this is equivalent to minimizing *q*(*x*), defined from the RNN dynamics of Eq. (1):

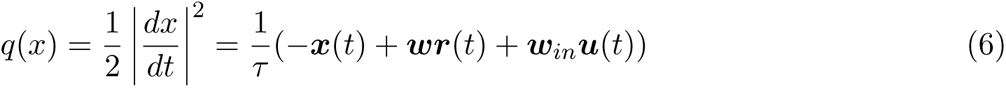

We repeated this search for every set of external inputs *u*(*t*), which included null inputs for late instruction trials, task cue (pro-DMS vs. anti-DMS), and attention cues (for the two-modality DMS).

For each optimization run, we randomly selected 200 out of the 1000 simulated trials to create initial conditions for the optimizer. From each trial, one initial condition was generated by taking the 1000-dimensional system state at the end of the first instruction period. Each optimization run had the following parameters: number of iterations = 30000; initial learning rate = 0.1; outlier distance scale = 10; unique tolerance = 10. The outlier distance scale servesd as a cutoff distance for discarding outlier putative fixed points, measured from the centroid of the vector of initial conditions. The unique tolerance is the numerical precision used to determine if two fixed points are unique. Fixed points that were too close to previously identified ones were discarded. Two fixed points were considered unique if their distance exceeded this threshold.

To visualize the fixed points and RNN neural trajectories, we plotted them in principal component space spanned by the top three principal components extracted from the average activities during the first instruction period. In addition, we visualized the entire landscape of the energy function *q*(*x*) in the top two principal components during the first instruction period along with their respective energy minima for each external input *u*(*t*) condition (Fig. 2a,b; Fig. 5b).

### CEBRA analysis

To visualize network dynamics and analyze the impact of top-down signaling on RNN dynamics, we applied unsupervised manifold discovery CEBRA [63] on the 1,000 simulated trials of representative RNN models during the first instruction and first stimulus periods. This method is capable of uncovering latent structures in multidimensional dynamical datasets, either unsupervised or conditioned on specific variables of interest. Throughout this study, we conducted unsupervised latent structure discovery based solely on the temporal sequence of data points across the entire dataset, without considering trial conditions, tasks, or stimulus identities. More specifically, we utilized the default model parameters for the ‘offset10-model’ architecture, using a batch size of 256, time offsets of 10, learning rates of 3*e −* 4, and cosine as the distance measure.

### Cross-temporal discriminability analysis

In addition to the SVM decoding analysis (see above), we also quantified the amount of information encoded by each unit within RNNs using cross-temporal discriminability analysis ([79–81]). For Figure 2j–m, the firing rate estimate time courses of each unit for each first stimulus identity were initially divided into two splits (odd vs. even trials) and averaged across trials within each split. Subsequently, for each unit in each split, the difference in the average firing rates between the two conditions based on the first stimulus identity (i.e., +1 and -1) was computed. Next, Pearson’s correlation coefficient between the two splits was calculated. The discriminability score was then obtained by applying Fisher’s *z*-transformation. This process was carried out separately for the attended modality (Fig. 2j,k) and the unattended modality (Fig. 2l,m). For Figure 7f, the first stimulus identity was fixed to -1, and the discriminability of the task cue signal was computed using the two task cue identities (pro- vs. anti-DMS). The diagonal values of the discriminability matrices are shown. For Figure 8d–f, similar procedures to those employed for Figure 2j–m were implemented.

### Centrality measure

We utilized a functional closeness centrality measure to determine the centrality of each unit within RNNs. For this, we first defined a functional distance metric *d_i,j_* between unit *i* and *j* a network, based on the absolute of the Pearson correlation (*ρ*) between their synaptic current activity over time, denoted as *x_i_* and *x_j_*:

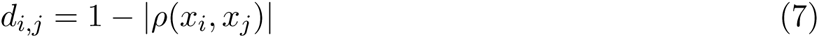

This function ensures that unit pairs that are more predictive of each other’s activity, either via positive or negative correlation, have a distance function closer to 0, while unit pairs with weak correlations have a distance function closer to 1.

Next, we used the complete distance matrix as the edge distances between nodes in a graph and employed the Python package *networkx* [82] to compute the network closeness centrality measure *C_i_* for each unit *i* from all *n* units:

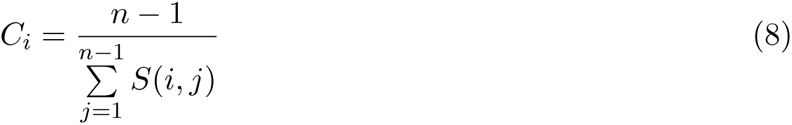

Here, *S*(*i, j*) represents the shortest path distance between two units *i* and *j*, computed utilizing the distance matrix *d_i,j_* via Dijkstra’s algorithm with the distance attributes as edge weights.

### Identifying task and attention cue selective units

To identify unit selective for pro-DMS vs. anti-DMS task signals, we first generated 400 trials for the one-modality DMS and 1,000 trials for the two-modality DMS. We next extracted firing rate estimates of all units in each of the trained RNNs. Then, the trials and firing rate estimates were sorted based on the early task cue signal, separating them into pro-DMS and anti-DMS groups. For each unit, we averaged the firing rate estimates during the first instruction window for each trial. Subsequently, we performed the one-sided Wilcoxon rank-sum statistical test on the average firing rates from the two task groups for each unit. Units with significantly higher average firing rates in the pro-DMS group compared to the anti-DMS group were classified as pro-DMS selective units. Similar procedures were performed to identify attention-selective units for the two-modality RNNs.

### Lesioning analyses

For Figure 4 and Figure 6, trained synaptic connections were lesioned systematically to identify specific connections important for encoding and maintenance of top-down information. For each trained network, connections were initially classified into four subgroups: *I → I* (inhibitory units inhibiting other inhibitory units), *I → E* (inhibitory units inhibiting excitatory units), *E → I* (excitatory units exciting inhibitory units), and *E → E* (excitatory units exciting other excitatory units). Within each subgroup, connections were further categorized based on their selectivity for the task and attention cue signal (see above). The selective connections were then lesioned by reducing their strength by 50%.

### Experimental data

To further characterize the impact of top-down signaling over information processing, we utilized publicly available experimental data on the role of recurrent processing in the activity of mouse primary visual cortex [54]. In this task, mice had to determine the location of a salient grating visual pattern which contrasted from a visual background, distinguishing them based either on the orientation, the phase, or the texture of the grating pattern. For the purposes of our analyses, we averaged results across orientation, phase, and texture conditions. Target patterns could appear either in the right or left visual field, prompting mice to indicate their locations by licking a spout on the corresponding side to receive water or milk droplets as rewards. Crucially, neural recordings and stimuli were set up in such a way that image contents falling within the receptive fields of the recorded neurons were strictly identical in the figure and background conditions. Thus, any difference in neural activity measured across conditions was caused by the context indicated by image regions outside of the measured neural receptive field.

We first sought to determine whether V1 activity contained information distinguishing target grated stimuli from non-target grated backgrounds even when the sensory input within receptive fields was identical. This would indicate that feedback from regions outside receptive fields into the measured V1 sites. For this analysis, neural activity consisted of multiunit activity from 198 recording sites in 13 different V1 electrode penetrations in 6 awake mice. To determine which information was contained in V1 multiunit activity, we performed a 4-fold cross-validated linear SVM decoding analysis (MATLAB function fitcsvm), for figure versus background labels from 0.5 s before to 1 s after stimulus presentation, utilizing activity in separate channels as features. To obtain a null distribution for decoding accuracy, we generated 1,000 random label permutations and performed the same decoding analysis.

Furthermore, we leveraged an additional analysis using the aforementioned dataset. Specifically, we determined whether activity of inhibitory neurons in V1 is necessary for figure versus background discrimination, as predicted by the central role of inhibitory neurons in our modeling approach. The analysis contained multiunit activities of different subgroups of inhibitory interneurons in V1: vasoactive intestinal peptide–expressing (VIP) and somatostatin-positive interneurons (SST). In addition, the dataset also contains recordings when VIP or SST neurons were optogenetically inhibited (see [54] for more details).

Using this dataset, we decoded figure versus background labels from V1 multiunit activity with a 4-fold cross-validated linear SVM, both with and without optogenetic VIP suppression. We then compared decoding accuracy between the laser-on and laser-off conditions by performing a two-sample one-sided Kolmogorov-Smirnov (KS) test for whether accuracy was larger in trials without laser suppression. We performed KS tests within the laser delivery time window (250 ms to 500 ms) and Bonferroni corrected p-values across different time bins.

### Statistical analyses

All RNNs trained in the present study were randomly initialized (with random seeds) before training. Nonparametric statistical methods were employed throughout the study. In figures containing boxplots, two-sided Wilcoxon rank-sum or signed-rank tests were performed to ascertain statistically significant differences between two groups.

Following linear SVM decoding and functional centrality analyses, we aimed to determine whether decoding accuracy or functional centrality values were higher in excitatory or inhibitory neurons. To achieve this, we utilized the scipy.stats function kstest and performed two-sided Kolmogorov-Smirnov (KS) tests across all time bins and Bonferroni corrected for the number of tested time bins. Similarly, we performed Bonferroni corrected one-sided KS tests across time to specifically test the hypothesis that decoding accuracies in V1 would decrease following optogenetical silencing.

For cross-temporal discriminability analyses, we performed binomial tests and Bonferroni corrected for the number of tests to determine whether each decoding accuracy value was higher than chance. Assuming a false positive rate of 5%, and a Bernoulli process to generate significant decoding events at this rate, the number of instances *S* falsely classified as significant within a population of size *N* is given by a binomial distribution: *S ∼ binomial*(*N,* 0.05). Accordingly, we derived a binomial p-value for the probability of obtaining an observed sensitive instance count *K* larger than expected by chance, using the cumulative binomial distribution: *p* = 1*−binomialCDF* (*K, N,* 0.05).

## Code availability

The code for the analyses performed in this study will be made available upon acceptance for publication.

## Data availability

The trained RNN models used in the present study will be deposited as MATLAB-formatted data in Open Science Framework upon acceptance for publication.

## Extended Data Figures

**Extended Data Fig. 1.**
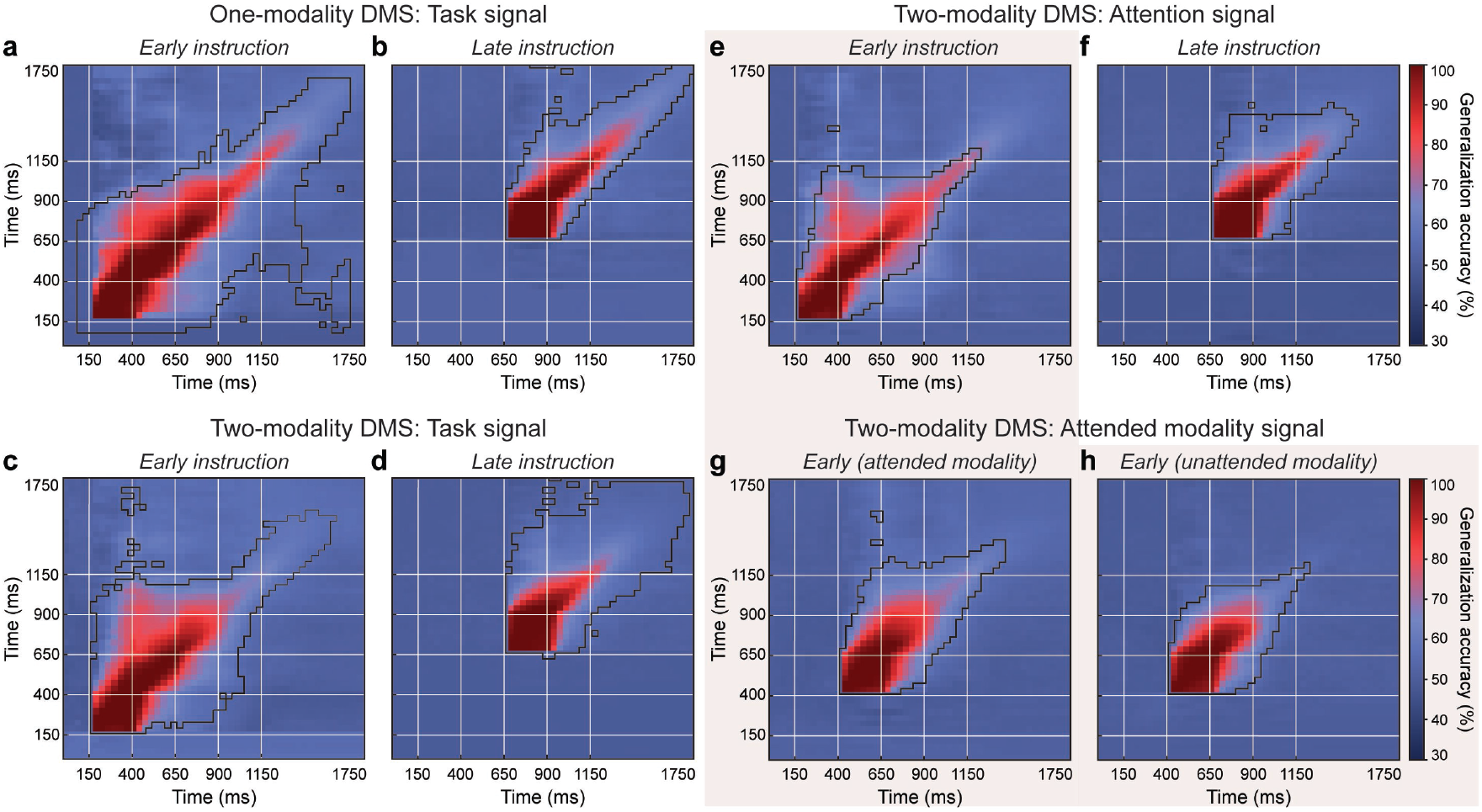
Temporal encoding of top-down signals during DMS tasks. Cross-temporal decodability of task and attention signals is shown as the average generalization accuracy for each pair of training/testing time windows. Highlighted boundaries indicate significant decoding accuracy, *p <* 1*e −* 20, binomial test, Bonferroni corrected. White lines denote the reference event of each trial period (task and attention cues, stimulus onsets, response). **a.** One-modality DMS, task signal, early instruction. **b.** Same, for late instruction. **c.** Two-modality DMS, task signal, early instruction. **d.** Same, for late instruction. **c.** Two-modality DMS, attention signal, early instruction **f.** Same, for late instruction. **g.** Two-modality DMS, attended modality signal, early instruction, attended modality. **h.** Same, for unattended modality. **i.** One-modality DMS, stimulus identity. **j.** Two-modality DMS, stimulus identity, stimulus modality 1. **k.** Same, for stimulus modality 2.

**Extended Data Fig. 2.**
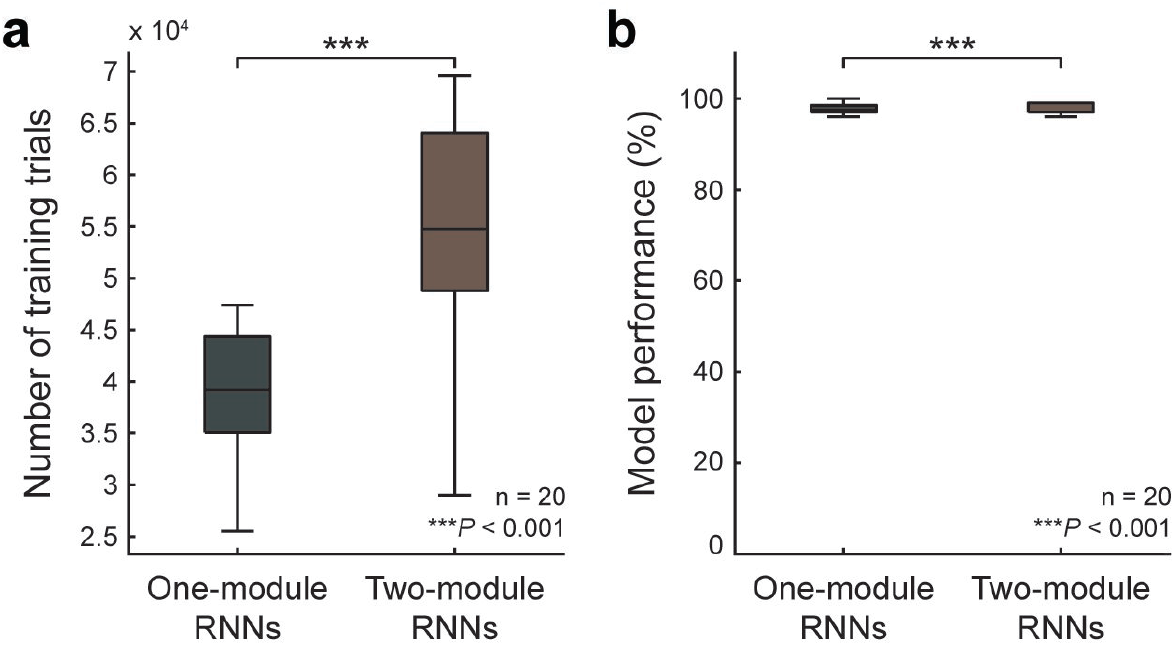
Training two-module RNNs takes longer for the one-modality DMS compared to the one-module RNNs. **a.** Comparison of the number of training trials required for training between the one-module and two-module RNNs, with 20 RNNs trained for each case. **b.** Both one-module and two-module RNNs achieved high accuracy in performing the one-modality DMS task (mean*±*stdev; 97.7*±*1.3% for the one-module model; 97.6*±*1.2% for the two-module model). Boxplot: central lines, median; bottom and top edges, lower and upper quartiles; whiskers, 1.5 × interquartile range; outliers are not plotted. \*\*\**p <* 0.001 by two-sided Wilcoxon rank-sum test.

**Extended Data Fig. 3.**
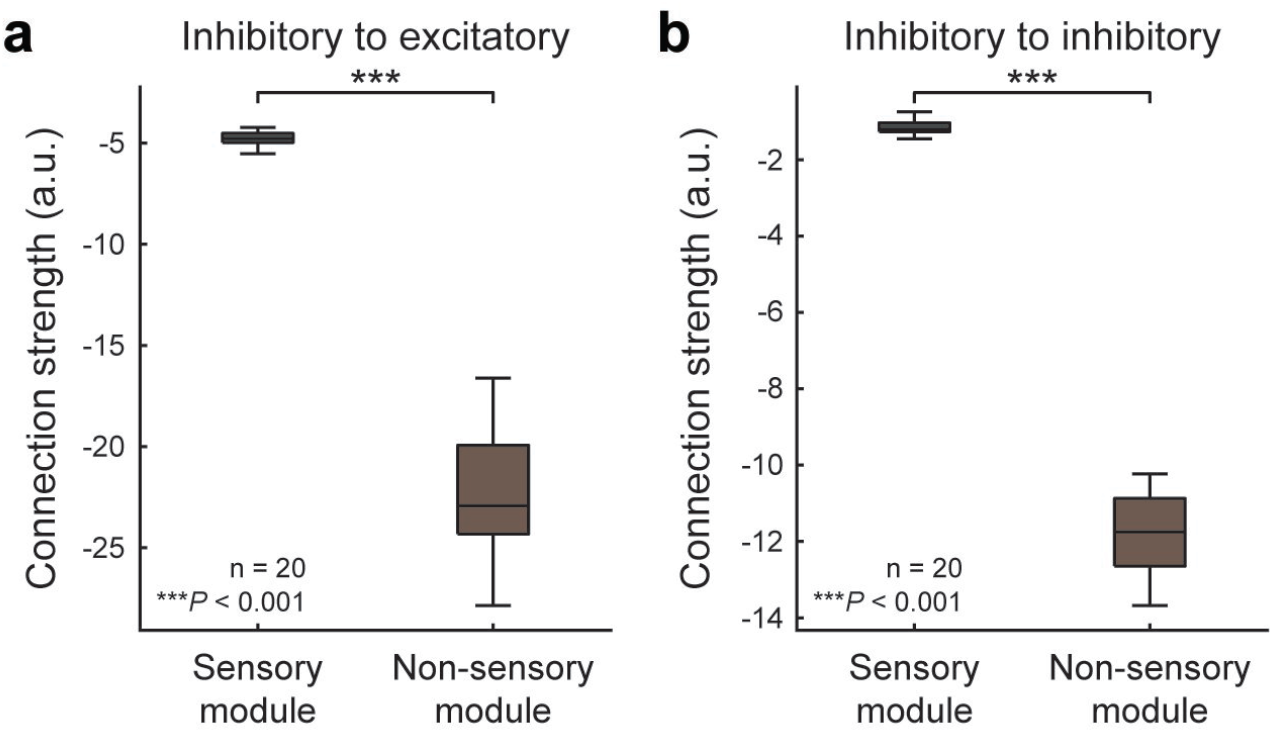
Comparison between *I → E* and *I → I* connection strength in two-module RNNs. *I → E* (**a**) and *I → I* (**b**) connections were significantly stronger within the non-sensory module than the sensory module. Boxplot: central lines, median; bottom and top edges, lower and upper quartiles; whiskers, 1.5 × interquartile range; outliers are not plotted. \*\*\**p <* 0.001 by two-sided Wilcoxon rank-sum test.

**Extended Data Fig. 4.**
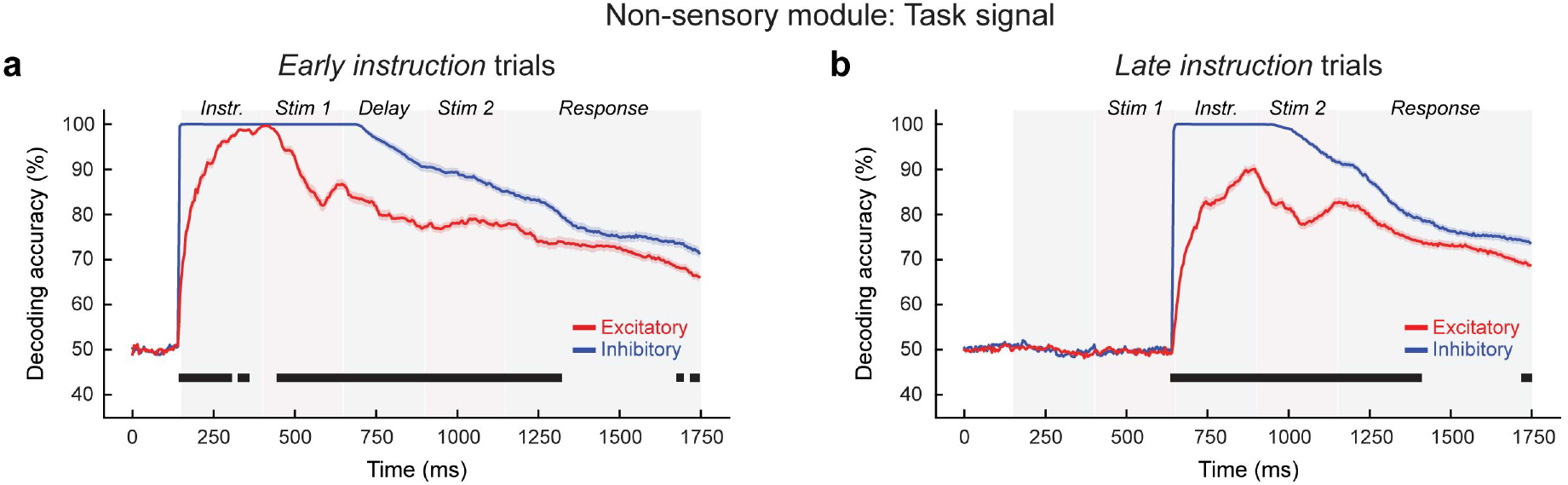
Decoding task signals in non-sensory module of two-module RNNs. **a.** Average decoding accuracy in early instruction trials for pro- vs. anti-DMS signal in the non-sensory module of two-module networks, including either excitatory (red) or inhibitory (blue) units. Shaded areas indicate standard error of the mean. Black bars indicate significant differences between excitatory and inhibitory units (*p <* 0.05, two-sided KS test). **b.** Same, for late instruction trials.

## References

1. Grossberg, S. & Grossberg, S. How does a brain build a cognitive code? Studies of mind and brain: neural principles of learning, perception, development, cognition, and motor control, 1–52 (1982).

2. Friston, K. A theory of cortical responses. Philosophical transactions of the royal society b: biological sciences 360, 815–836 (2005).

3. Mumford, D. On the computational architecture of the neocortex: ii the role of cortico-cortical loops. Biological cybernetics 66, 241–251 (1992).

4. Fenske, M. J., Aminoff, E., Gronau, N. & Bar, M. Top-down facilitation of visual object recognition: object-based and context-based contributions. Progress in brain research 155, 3–21 (2006).

5. Bar, M. et al. Top-down facilitation of visual recognition. Proceedings of the national academy of sciences 103, 449–454 (2006).

6. Bullier, J. Integrated model of visual processing. Brain research reviews 36, 96–107 (2001).

7. Rungratsameetaweemana, N., Squire, L. R. & Serences, J. T. Preserved capacity for learning statistical regularities and directing selective attention after hippocampal lesions. Proceedings of the national academy of sciences 116, 19705–19710 (2019).

8. Melloni, L., van Leeuwen, S., Alink, A. & Müller, N. G. Interaction between bottom-up saliency and top-down control: how saliency maps are created in the human brain. Cerebral cortex 22, 2943–2952 (2012).

9. Mante, V., Sussillo, D., Shenoy, K. V. & Newsome, W. T. Context-dependent computation by recurrent dynamics in prefrontal cortex. Nature 503, 78–84 (2013).

10. Cole, M. W. et al. Multi-task connectivity reveals flexible hubs for adaptive task control. Nature neuroscience 16, 1348–1355 (2013).

11. Remington, E. D., Narain, D., Hosseini, E. A. & Jazayeri, M. Flexible sensorimotor computations through rapid reconfiguration of cortical dynamics. Neuron 98, 1005–1019 (2018).

12. Schapiro, A. C., Gregory, E., Landau, B., McCloskey, M. & Turk-Browne, N. B. The necessity of the medial temporal lobe for statistical learning. Journal of cognitive neuroscience 26, 1736–1747 (2014).

13. Uddin, L. Q. Cognitive and behavioural flexibility: neural mechanisms and clinical considerations. Nature reviews neuroscience 22, 167–179 (2021).

14. Douw, L., Wakeman, D. G., Tanaka, N., Liu, H. & Stufflebeam, S. M. State-dependent variability of dynamic functional connectivity between frontoparietal and default networks relates to cognitive flexibility. Neuroscience 339, 12–21 (2016).

15. Voss, J. L., Gonsalves, B. D., Federmeier, K. D., Tranel, D. & Cohen, N. J. Hippocampal brain-network coordination during volitional exploratory behavior enhances learning. Nature neuroscience 14, 115–120 (2011).

16. Luck, S. J., Chelazzi, L., Hillyard, S. A. & Desimone, R. Neural mechanisms of spatial selective attention in areas v1, v2, and v4 of macaque visual cortex. Journal of neurophysiology 77, 24–42 (1997).

17. Zhang, S. et al. Long-range and local circuits for top-down modulation of visual cortex processing. Science 345, 660–665 (2014).

18. Morales-Gregorio, A. et al. Neural manifolds in v1 change with top-down signals from v4 targeting the foveal region. Biorxiv, 2023–06 (2023).

19. McBride, E. G., Lee, S.-Y. J. & Callaway, E. M. Local and global influences of visual spatial selection and locomotion in mouse primary visual cortex. Current biology 29, 1592–1605 (2019).

20. Saenz, M., Buracas, G. T. & Boynton, G. M. Global effects of feature-based attention in human visual cortex. Nature neuroscience 5, 631–632 (2002).

21. Serences, J. T. & Boynton, G. M. Feature-based attentional modulations in the absence of direct visual stimulation. Neuron 55, 301–312 (2007).

22. Itthipuripat, S., Garcia, J. O., Rungratsameetaweemana, N., Sprague, T. C. & Serences, J. T. Changing the spatial scope of attention alters patterns of neural gain in human cortex. Journal of neuroscience 34, 112–123 (2014).

23. Debes, S. R. & Dragoi, V. Suppressing feedback signals to visual cortex abolishes attentional modulation. Science 379, 468–473 (2023).

24. Liu, T., Larsson, J. & Carrasco, M. Feature-based attention modulates orientation-selective responses in human visual cortex. Neuron 55, 313–323 (2007).

25. Henderson, M. M., Serences, J. T. & Rungratsameetaweemana, N. Categorization dynamically alters representations in human visual cortex. Biorxiv (2023).

26. Rademaker, R. L., Chunharas, C. & Serences, J. T. Coexisting representations of sensory and mnemonic information in human visual cortex. Nature neuroscience 22, 1336–1344 (2019).

27. Musall, S., Kaufman, M. T., Juavinett, A. L., Gluf, S. & Churchland, A. K. Single-trial neural dynamics are dominated by richly varied movements. Nature neuroscience 22, 1677–1686 (2019).

28. Lab, I. B. et al. A brain-wide map of neural activity during complex behaviour. Biorxiv, 2023– 07 (2023).

29. Ebrahimi, S. et al. Emergent reliability in sensory cortical coding and inter-area communication. Nature 605, 713–721 (2022).

30. Alreja, A., Nemenman, I. & Rozell, C. J. Constrained brain volume in an efficient coding model explains the fraction of excitatory and inhibitory neurons in sensory cortices. Plos computational biology 18, e1009642 (2022).

31. Melzer, S. & Monyer, H. Diversity and function of corticopetal and corticofugal gabaergic projection neurons. Nature reviews neuroscience 21, 499–515 (2020).

32. Schroeder, A. et al. Inhibitory top-down projections from zona incerta mediate neocortical memory. Neuron 111, 727–738 (2023).

33. Pfeffer, C. K., Xue, M., He, M., Huang, Z. J. & Scanziani, M. Inhibition of inhibition in visual cortex: the logic of connections between molecularly distinct interneurons. Nature neuroscience 16, 1068–1076 (2013).

34. Pi, H.-J. et al. Cortical interneurons that specialize in disinhibitory control. Nature 503, 521– 524 (2013).

35. Herstel, L. J. & Wierenga, C. J. Network control through coordinated inhibition. Current opinion in neurobiology 67, 34–41 (2021).

36. Isaacson, J. S. & Scanziani, M. How inhibition shapes cortical activity. Neuron 72, 231–243 (2011).

37. Keller, A. J. et al. A disinhibitory circuit for contextual modulation in primary visual cortex. Neuron 108 (2020).

38. Wilmes, K. A. & Clopath, C. Inhibitory microcircuits for top-down plasticity of sensory representations. Nature communications 10, 5055 (2019).

39. Lehr, A. B., Kumar, A. & Tetzlaff, C. Sparse clustered inhibition projects sequential activity onto unique neural subspaces. Biorxiv, 2023–09 (2023).

40. Shen, B., Louie, K. & Glimcher, P. W. Flexible control of representational dynamics in a disinhibition-based model of decision making. Elife 12, e82426 (2023).

41. Panzeri, S., Brunel, N., Logothetis, N. K. & Kayser, C. Sensory neural codes using multiplexed temporal scales. Trends in neurosciences 33, 111–120 (2010).

42. Gire, D. H., Whitesell, J. D., Doucette, W. & Restrepo, D. Information for decision-making and stimulus identification is multiplexed in sensory cortex. Nature neuroscience 16, 991–993 (2013).

43. Song, H. F., Yang, G. R. & Wang, X.-J. Training excitatory-inhibitory recurrent neural networks for cognitive tasks: a simple and flexible framework. Plos computational biology 12, e1004792 (2016).

44. Miconi, T. Biologically plausible learning in recurrent neural networks reproduces neural dynamics observed during cognitive tasks. Elife 6, e20899 (2017).

45. Yang, G. R., Joglekar, M. R., Song, H. F., Newsome, W. T. & Wang, X.-J. Task representations in neural networks trained to perform many cognitive tasks. Nature neuroscience 22, 297–306 (2019).

46. Barbosa, J. et al. Early selection of task-relevant features through population gating. Nature communications 14, 6837 (2023).

47. Park, I. M., Ságodi, Á. & Sokół, P. A. Persistent learning signals and working memory without continuous attractors. Arxiv preprint arxiv:2308.12585 (2023).

48. Roach, J. P., Churchland, A. K. & Engel, T. A. Choice selective inhibition drives stability and competition in decision circuits. Nature communications 14, 147 (2023).

49. Kim, R., Li, Y. & Sejnowski, T. J. Simple framework for constructing functional spiking recurrent neural networks. Proceedings of the national academy of sciences 116, 22811–22820 (2019).

50. Kleinman, M., Chandrasekaran, C. & Kao, J. C. Recurrent neural network models of multi-area computation underlying decision-making. Biorxiv, 798553 (2019).

51. Michaels, J. A., Schaffelhofer, S., Agudelo-Toro, A. & Scherberger, H. A goal-driven modular neural network predicts parietofrontal neural dynamics during grasping. Proceedings of the national academy of sciences 117, 32124–32135 (2020).

52. Kim, R. & Sejnowski, T. J. Strong inhibitory signaling underlies stable temporal dynamics and working memory in spiking neural networks. Nature neuroscience 24, 129–139 (2021).

53. Kim, Y. et al. Brain-wide maps reveal stereotyped cell-type-based cortical architecture and subcortical sexual dimorphism. Cell 171, 456–469.e22 (2017).

54. Kirchberger, L. et al. The essential role of recurrent processing for figure-ground perception in mice. Science advances 7, eabe1833 (2021).

55. Blough, D. S. Delayed matching in the pigeon. Journal of the experimental analysis of behavior 2, 151 (1959).

56. Hampson, R. E., Heyser, C. J. & Deadwyler, S. A. Hippocampal cell firing correlates of delayed-match-to-sample performance in the rat. Behavioral neuroscience 107, 715 (1993).

57. Mishkin, M. & Delacour, J. An analysis of short-term visual memory in the monkey. Journal of experimental psychology: animal behavior processes 1, 326 (1975).

58. Wu, Z. et al. Context-dependent decision making in a premotor circuit. Neuron 106, 316–328 (2020).

59. Monk, C. S. et al. Human hippocampal activation in the delayed matching-and nonmatching-to-sample memory tasks: an event-related functional mri approach. Behavioral neuroscience 116, 716 (2002).

60. Sussillo, D. & Barak, O. Opening the black box: low-dimensional dynamics in high-dimensional recurrent neural networks. Neural computation 25, 626–649 (2013).

61. Golub, M. D. & Sussillo, D. Fixedpointfinder: a tensorflow toolbox for identifying and characterizing fixed points in recurrent neural networks. Journal of open source software 3, 1003 (2018).

62. Driscoll, L., Shenoy, K. & Sussillo, D. Flexible multitask computation in recurrent networks utilizes shared dynamical motifs. Biorxiv, 2022–08 (2022).

63. Schneider, S., Lee, J. H. & Mathis, M. W. Learnable latent embeddings for joint behavioural and neural analysis. Nature, 1–9 (2023).

64. Bastos, G. et al. Top-down input modulates visual context processing through an interneuron-specific circuit. Cell reports 42 (2023).

65. Chini, M., Pfeffer, T. & Hanganu-Opatz, I. An increase of inhibition drives the developmental decorrelation of neural activity. Elife 11 (eds de la Prida, L. M., Colgin, L. L. & Vanhatalo, S.) e78811 (2022).

66. Fino, E. & Yuste, R. Dense inhibitory connectivity in neocortex. Neuron 69, 1188–1203 (2011).

67. Fu, Y. et al. A cortical circuit for gain control by behavioral state. Cell 156, 1139–1152 (2014).

68. Karnani, M. M. et al. Opening holes in the blanket of inhibition: localized lateral disinhibition by vip interneurons. Journal of neuroscience 36, 3471–3480 (2016).

69. Xu, H. et al. A disinhibitory microcircuit mediates conditioned social fear in the prefrontal cortex. Neuron 102, 668–682 (2019).

70. Letzkus, J. J. et al. A disinhibitory microcircuit for associative fear learning in the auditory cortex. Nature 480, 331–335 (2011).

71. Lee, S., Kruglikov, I., Huang, Z. J., Fishell, G. & Rudy, B. A disinhibitory circuit mediates motor integration in the somatosensory cortex. Nature neuroscience 16, 1662–1670 (2013).

72. Wang, X.-J. & Yang, G. R. A disinhibitory circuit motif and flexible information routing in the brain. Current opinion in neurobiology 49, 75–83 (2018).

73. Werbos, P. J. Backpropagation through time: what it does and how to do it. Proceedings of the ieee 78, 1550–1560 (1990).

74. Rungratsameetaweemana, N., Kim, R. & Sejnowski, T. J. Internal noise promotes heterogeneous synaptic dynamics important for working memory in recurrent neural networks. Biorxiv. eprint: https://www.biorxiv.org/content/early/2022/10/18/2022.10.14.512301.full.pdf (2022).

75. Hendry, S. H., Schwark, H., Jones, E. & Yan, J. Numbers and proportions of gaba-immunoreactive neurons in different areas of monkey cerebral cortex. Journal of neuroscience 7, 1503–1519 (1987).

76. Sherwood, C. C. et al. Scaling of inhibitory interneurons in areas v1 and v2 of anthropoid primates as revealed by calcium-binding protein immunohistochemistry. Brain, behavior and evolution 69, 176–195 (2007).

77. Duarte, R., Seeholzer, A., Zilles, K. & Morrison, A. Synaptic patterning and the timescales of cortical dynamics. Current opinion in neurobiology 43, 156–165 (2017).

78. Zador, A. M. & Dobrunz, L. E. Dynamic synapses in the cortex. Neuron 19, 1–4 (1997).

79. Wasmuht, D. F., Spaak, E., Buschman, T. J., Miller, E. K. & Stokes, M. G. Intrinsic neuronal dynamics predict distinct functional roles during working memory. Nature communications 9, 3499 (2018).

80. Spaak, E., Watanabe, K., Funahashi, S. & Stokes, M. G. Stable and dynamic coding for working memory in primate prefrontal cortex. Journal of neuroscience 37, 6503–6516 (2017).

81. Stokes, M. G. et al. Dynamic coding for cognitive control in prefrontal cortex. Neuron 78, 364–375 (2013).

82. Hagberg, A., Swart, P. & S Chult, D. Exploring network structure, dynamics, and function using networkx tech. rep. (Los Alamos National Lab.(LANL), Los Alamos, NM (United States), 2008).

